# Functionally Distinct pTfh1 Subsets with Roles in Malaria-Specific Immunity

**DOI:** 10.1101/2025.02.27.640479

**Authors:** Megan SF Soon, Damian A Oyong, Nicholas Dooley, Reena Mukhiya, Zuleima Pava, Dean Andrew, Jessica R Loughland, James McCarthy, Jo-Anne Chan, James G Beeson, Christian Engwerda, Ashraful Haque, Michelle J Boyle

## Abstract

Antibodies induced by infection or vaccination are essential mediators of protection. Induction of these protective responses is mediated by T-follicular CD4 T (Tfh) cells, and targeting these cells may be a strategy to boost antibody mediate protection. In humans, Tfh cells are analysed based on expression of CXCR3 and CCR6, with different subsets of Tfh (Tfh1, Tfh2, Tfh17) associated with antibody induction in a context-dependent manner. Here we dissected Tfh cells heterogeneity in healthy donors and individuals during controlled human malaria infection using scRNAseq. We identified two distinct Tfh1-like subsets with functional relevance, defined based on CCR7 expression. CCR7^neg^ Tfh1 cells express markers of cytotoxicity, while CCR7^pos^ Tfh1 cells produce reduced inflammatory cytokines and similar IL-21 resulting in unique cytokine milieu. In controlled human malaria infection, both CCR7^pos^ and CCR7^neg^ Tfh1, along with Tfh2 cells, clonally expanded and were transcriptionally and phenotypically activated. However, only CCR7^pos^ Tfh1 and Tfh2 cells associated with antibody development, suggesting a role for both these Tfh subsets in promoting humoral immunity to malaria. Data identify specific protective Tfh subsets that can be targeted to improve antibody mediated protection to malaria induced, and provide a frame work to dissect the role of Tfh subsets in other disease contexts.

## Introduction

Antibodies and B cells induced by infection or vaccination are essential mediators of protection from most pathogens. This includes malaria, caused by *Plasmodium falciparum* infection, which remains one of the largest killers globally with >200 million cases and 600 000 deaths annually ^1^. While the introduction of two malaria vaccines, RTS,S/AS01 and R21/Matrix-M, bring new hope to malaria control efforts, these vaccines only have modest and variable efficacy that is short-lived ^2,3^. Antibody development is driven by T-follicular helper (Tfh) cells, the key CD4 T cell subset that function within the germinal centre to support B cell activation. Cytokine and receptor mediated signals from Tfh cells, drive B cell selection, support antibody class switching, and differentiation into antibody-producing long-lived plasma cells and memory B cells. As such, Tfh cells are attractive targets to boost naturally acquired or vaccine-mediated protection and longevity ^4^. To develop such therapies, an improved understanding of how Tfh cells drive antibody development in malaria and specific disease contexts is required.

Tfh cells, identified by expression of C-X-C chemokine receptor 5 (CXCR5) and programmed cell death protein (PD1), are recognised to exist in distinct subsets, which have specific and context dependent roles in antibody induction. These Tfh subsets take on characteristics of CD4+ T cell helper cell lineages ^5^, and are commonly analysed by CXCR3 and CCR6 expression to identify Tfh1 (CXCR3+/CCR6-), Tfh2 (CXCR3-/CCR6-) and Tfh17 (CXCR3-/CCR6+) cells, both in the periphery and also in (GC) Tfh cells in secondary lymphoid tissues ^6^. The development and activation of Tfh subsets may be underpinned by ‘skewing’ of Tfh cells by the cytokine milieu induced during infection or vaccination, allowing Tfh cells to adopt specific roles to influence B cell development ^5,7^. Indeed, defined Tfh cell subsets have differing abilities to activate naïve or memory B cells and have been associated with antibody development in a pathogen and vaccine-dependent manner. For example, Tfh2 cells have the highest capacity to activate naïve B cells, and peripheral (p)Tfh2 cells have been associated with the development of broadly neutralizing antibodies in HIV infection ^8^. On the other hand, Tfh1 cells robustly activate memory B cells, and the activation of pTfh1 cells are consistently associated with antibodies induced by influenza vaccination ^9,10^. Thus, development of strategies to target Tfh cells to boost antibody development require a context specific and in-depth understanding of specialised Tfh subsets that drive protective antibodies.

In malaria, progress has been made to define roles of Tfh cells in antibody development in infection and vaccination ^11^. Studies by our team have shown that the activation of pTfh2 cells was associated with antibody development during controlled human malaria infection (CHMI) with *P. falciparum* ^12^. Similarly, others have shown that pTfh2 cells are associated with induction of antibodies by approved and experimental malaria vaccines ^13–15^. These association are observed despite the overall response during malaria being dominated by pTfh1 cells ^12,16,17^. Further, in Malian children, pTfh1 cell activation was not associated with antibody development following malaria treatment ^17^, and pTfh1 cells have been associated with induction of short-lived plasma cells which may hamper the germinal centre ^12,18^. As such, data suggests that strategies that can boost Tfh2 and reduce Tfh1 activation may improve malaria antibody development. However, recent studies by our team have shown that these relationships are more nuanced, with Ugandan children who had the highest levels of protective antibodies having increased frequencies of activated pTfh2 and pTfh1 cells ^19^. Additionally, parasite activated pTfh1 cells induced T-bet+ B cells with high IgG3 expression ^20^, and IgG3 antibodies are consistently associated with protection from malaria ^21^. To develop Tfh cell targeted approaches to improve malaria antibody development and longevity in infection and vaccination, an increased understanding of the specific Tfh cells that drive protective antibodies is required.

We hypothesised that contradictory results may be in part due to the limited approaches to identify subsets of Tfh cells, and an improved understanding of Tfh heterogeneity would reveal protective subsets associated with antibody development in malaria. In this study, we applied scRNAseq and spectral flow cytometry to unique clinical cohorts to discover functionally relevant Tfh cell subsets that have differing roles in malaria antibody development. We identify two transcriptionally and phenotypically distinct pTfh1 subsets in healthy individuals which can be identified based on CCR7 expression, with differences in expression of cytotoxic markers and cytokine production. Tracking pTfh cell activation with sc-RNA/VDJ-seq during CHMI, we show Tfh subset specific activation and clonal expansion, with increased activation and clonal expansion of pTfh1 CCR7^pos^ compared to other subsets. Finally, we show that antigen specific pTfh1 CCR7^pos^ cells, along with pTfh2 cells, but not pTfh1 CCR7^neg^ cells associated with development of functional and protective antibodies following malaria. Our data explain previous contradictions in the field and highlight the phenotypic and functional diversity of Tfh cells during human infection. Data open new avenues to develop approaches which harness Tfh cells to enhances vaccine efficacy and improve antibody development and protection.

## Results

### Analysis of Tfh cells with scRNAseq identifies two Tfh1-like subsets

We hypothesised that hidden diversity of Tfh cells with distinct functional capacity, not captured in traditional analysis methods, underpinned the unclear roles of Tfh subset in antibody development in malaria. To dissect this diversity and identify protective Tfh subset that drive malaria antibody development we first performed scRNAseq on pTfh cells sorted from healthy human donors (n=4, sex 50% males, aged 28.5 [25.5-35], median [IQR] years). To understand the link between commonly identified Tfh cells and potential hidden diversity, we sorted three Tfh subsets (pTfh1, pTfh2 and pTfh17 from CXCR5+ PD1+ cells), based on classical definitions of CXCR3/CCR6 expression, alongside the less functional or less activated CXCR5+ PD1- compartment (**Fig. 1A, Supplementary Fig. S1A**). After QC, a total of 7,302 cells were analysed with high quality cells captured from each donor and subset (pTfh1 n= 2,089, pTfh2 n=717, pTfh17 n=744, PD1- n=2,839, **Supplementary** Fig. 1B). Following data integration by donor (**Supplementary** Fig. 1C), we identified 11 Tfh cell clusters using a combination of unsupervised and semi-supervised clustering methods (**Fig. 1B-D, Supplementary Table 1**). Strikingly, two distinct Tfh1 clusters were identified, both with relatively high expression of *CXCR3* and *CST7* (annotated Tfh1_cytotoxic and Tfh1_CCR7). A Tfh2 cluster was identified with high expression of genes associated with GC Tfh cells, including *TOX2* ^22,23^*, MAF* ^24,25^, and *TCF7*. The signature of this cluster mapped to the majority of tonsil GC Tfh cells in a publicly available data set ^26^ (**Supplementary** Fig. 1D), confirming the transcriptional relationship between Tfh2 cluster and GC Tfh cells identified here and reported previously ^8^. Tfh17 cluster was annotated based on high expression of genes associated with Th17 cells *CCR6, BHLHE40,* and *TGFB1* ^27–29^. Most cells within these Tfh1/Tfh2 and Tfh17 transcriptional clusters were sorted from the PD1+ cells (**Fig. 1B/C**). Other transcriptional clusters included cells enriched in Type I IFN genes (annotated as IFNI), a cluster enriched with heat shock proteins (annotated stress), another cluster enriched with ribosomal genes (annotated ribosomal), and two clusters without clear marker genes (cluster 2, possibly a quiescence cell subset, with upregulation of genes *BTG1* and *ZFP36L2,* and cluster 3 which only had limited upregulated genes including annexin family members *ANXA1*, *ANXA2*, *S100A10)* (**Fig. 1D**). Analysis of key T helper genes also identified a small cluster of cells positioned close to the Tfh1 cluster that expressed a mixture of Tfh1/Tfh17 signature, which were manually clustered and annotated as Tfh1.17 (**Fig. 1B/D**). Additionally, another subset of cells expressing *FOXP3* and Treg signature genes including *CTLA4* was identified manually and annotated as Tfr (**Fig. 1D**). The remaining Tfh cell subsets expressed diverse levels of markers associated with various aspects of Tfh or CD4 T cell development, with no clear patterns of expression across subsets (**Supplementary** Fig. 1E).

**Fig. 1.**
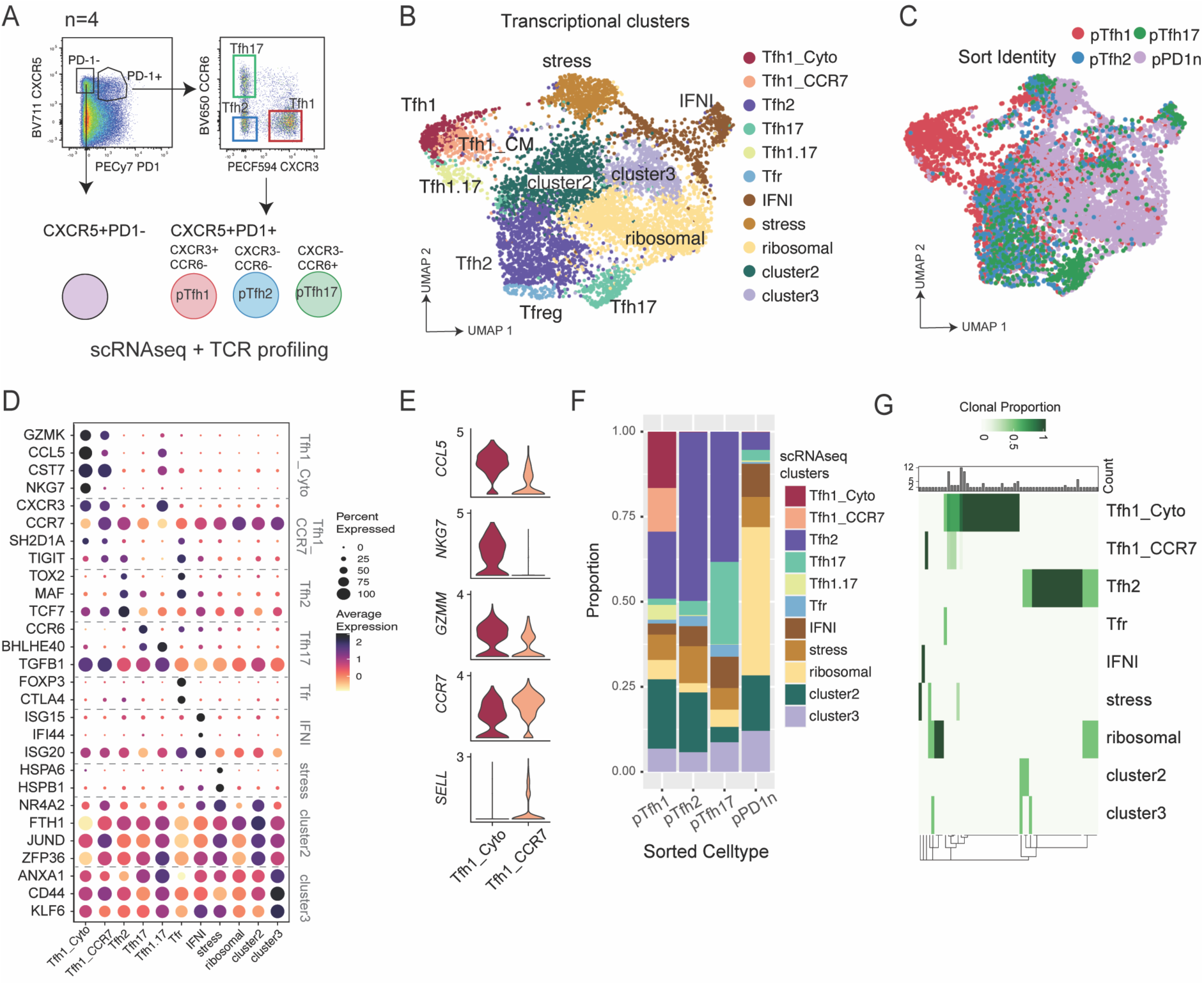
scRNAseq analysis of pTfh cells from healthy individuals. **A.** Four populations were sorted from pTfh CD4+ cells from 4 healthy donors - 1) CXCR5+PD1-, 2) CXCR5+PD1+ pTfh1, 3) CXCR5+PD1+ pTfh2, and 4) CXCR5+PD1+ pTfh17. Populations were analysed by 5’ droplet scRNAseq for gene expression and TCR sequence. **B.** UMAP of data following integration by donor, with 11 transcriptional subsets indicated. **C.** UMAP of data depicting cell sorted identity. **D.** Relative gene expression of genes used to annotate transcriptional clusters. **E.** Average expression of key genes in Tfh1_Cyto and Tfh1_CCR7 annotated transcriptional clusters. **F.** Relationship between cell sorted identity and transcriptional cluster identity in the data set. **G.** Fate sharing at the individual clonotype level across transcriptional subsets. Expanded clonotypes (n³2) with only same fates (exist within the same subset), or different fates are shown See also Supplementary Fig. 1 and Supplementary Table S1 and S2.

Comparing the two cell clusters identified as Tfh1 cells, the Tfh1_cytotoxic cluster had high expression of *GZMK*, *GZMM*, *CCL5* and *NKG7* (**Fig. 1D/E, Supplementary** Fig. 1E**, Supplementary Table 2)**. The signature of the Tfh1_cytotoxic annotated cells overlaps with cytotoxic Tfh cells previously identified in mice models, including the increased expression of *EOMES* ^30^, and in a recent small analysis of human Tfh cells from the periphery and tonsil ^31^ (**Supplementary** Fig. 1E). Potentially cytotoxic, granzyme-K expressing pTfh cells have also recently been identified in type-1 skewed response in mice, and in human CMV specific pTfh cells ^32^. In contrast, the Tfh1_CCR7 cluster had lower expression of cytotoxic markers, and instead had increased expression of *CCR7* and *SELL.* Differential expression of CCR7 along with PD1 has previously been used to differentiate Tfh subsets within CXCR5+ cells in both mice and humans, with CCR7^lo^/PD1^hi^ described as effector cells and CCR7^hi^/PD1^lo^ described as resting cells ^33^. However here, both Tfh1_cytotoxic and Tfh1_CCR7 subsets largely existed in clusters that were from the PD1+ sorted populations thus identifying an unappreciated gradient of CCR7 expression within PD1+ cells (**Fig. 1C/F, Supplementary** Fig. 1F). Pathway analysis of Tfh1_CCR7 and Tfh1_cytotoxic cluster marker genes highlighted the shared and divergent transcriptional signatures of these cells (**Supplementary** Fig. 1G). For example, both subsets were enriched for ‘Th1 pathway’, and ‘Interferon gamma signalling’, while only Tfh1_CCR7 cells were enriched in ‘Th2 pathway’ and ‘ICOS-ICOSL Signalling in T Helper Cells’ suggesting potential functional differences between the two transcriptional clusters (**Supplementary** Fig. 1G). Together with the distinct expression of cytotoxic markers, data suggests Tfh1 cells exist in two distinct and functionally relevant subests.

To dissect the relationship between the diversity of Tfh cells identified transcriptional, and the phenotypic subsets these cells would have been characterized as with flow cytometry, the degree of overlap between designated Tfh cell subsets and sorted cell phenotypes was analysed. Tfh1_cytotoxic, Tfh1_CCR7, Tfh2 and Tfh17 clusters, defined by transcriptional profiles, were mainly found in cells expressing *PDCD1*. Further, *CXCR3* and *CCR6* expression overlapped with the pTfh1 and Tfh17 sorted populations, respectively, suggesting some overlap between transcriptional clusters and those identified via CXCR3 and CCR6 expression (**Fig. 1C**). Nearly all cells found within both Tfh1 transcriptional clusters (Tfh1_cytotoxic and Tfh1_CCR7), were from the pTfh1 sorted cell population. However, the sorted pTfh1 population also contained cells from all other transcriptional subsets.

Similarly, the phenotypic pTfh2 population was dominated by the Tfh2 transcriptional subset but also contained cells with diverse transcriptional annotations (**Fig. 1F, Supplementary** Fig. 1F). This pattern was also seen for the phenotypic pTfh17 population. The PD1- phenotypic cells contained cells of all clusters, but were dominated by IFNI, stress, ribosomal and cluster 2 and cluster 3 cells. Thus, while commonly used flow cytometry strategies do overlap with transcriptional diversity, further phenotypic markers would be required to fully dissect Tfh cell subsets.

To understand the clonal sharing between clusters, TCR sequences were analysed. TCR sequences with complete TRA/TRB genes were captured for 3,998 (55.6%) cells. Clones were defined by the sharing of identical amino acid sequences. While the majority of TRA/TRB cells were singletons (3839 clones), clonal expansion was detected in 159 cells across 57 clones ranging in size from 2 to 20 identical clones. Clonal expansion was detected particularly within the Tfh1_cytotoxic subset (**Supplementary** Fig. 1H). As expected, no clones were shared across individuals (**Supplementary** Fig. 1I) Within each clonotype, we examined the fate sharing clones to investigate if clones of the same family tend to share the same transcriptional subsets or were found across diverse Tfh phenotypes. 40/57 clonotypes (70.2%) were found in a single subset. Clonal sharing was evident between Tfh1_cytotoxic and Tfh1_CCR7 but not Tfh2 transcriptional clusters, while Tfh2 shared clones with ribosomal, cluster 2 and cluster 3 cells (**Fig. 1G**). Together, these data show that while commonly used phenotypic markers CXCR3 and CCR6 can resolve some heterogeneity in pTfh cells, alternative distinct Tfh subsets, particularly within the pTfh1 compartment require additional phenotypic markers to dissect subsets of potential functional relevance.

### pTfh1 subsets with distinct phenotypic and functional characteristics can be defined by CCR7 expression

Transcriptional analysis identified two pTfh1-like subsets with distinct expression of cytotoxic and memory associated genes suggesting functional relevance. To assess if these subsets could be identified phenotypically and develop an approach to identify these subsets along with pTfh2 and pTfh17 cells in clinical cohorts, we designed a comprehensive phenotyping panel for pTfh cells which included commonly assessed chemokine receptors (CXCR3 and CCR6), markers identified in our transcriptional data set or with reported roles in Tfh cell function or development (**Supplementary Table S3)**. To test this approach to define Tfh subsets, we first analysed pTfh cells in healthy human donors (n= 19, sex 83.3% males, aged 24 years [20.2-28.2, median [IQR] years). pTfh cells were identified based on CXCR5 and PD1 expression, and then analysed with unsupervised clustering considering CXCR3, CCR6, CCR7, NKG7, GrzmB, GrzmK, cMAF, TIGIT and FoxP3. As seen in transcriptomic analysis, unbiased clustering consistently identified two subsets of pTfh1 cells based on CCR7 expression, where CCR7^neg^ cells also expressed high levels of NKG7, GrzmB and GrzmK, while CCR7^pos^ cells were negative for these markers (**Fig. 2A, Supplementary** Fig. 2A-C). pTfh2, pTfh17 and pTfh1-17 cells could also be identified with unbiased clustering (**Fig. 2A**). Additionally, FoxP3+ Tfr cells were identified and largely clustered within the pTfh2 cell subset, consistent with scRNAseq data (**Fig. 2A, Supplementary** Fig. 2A-C).

**Figure 2:**
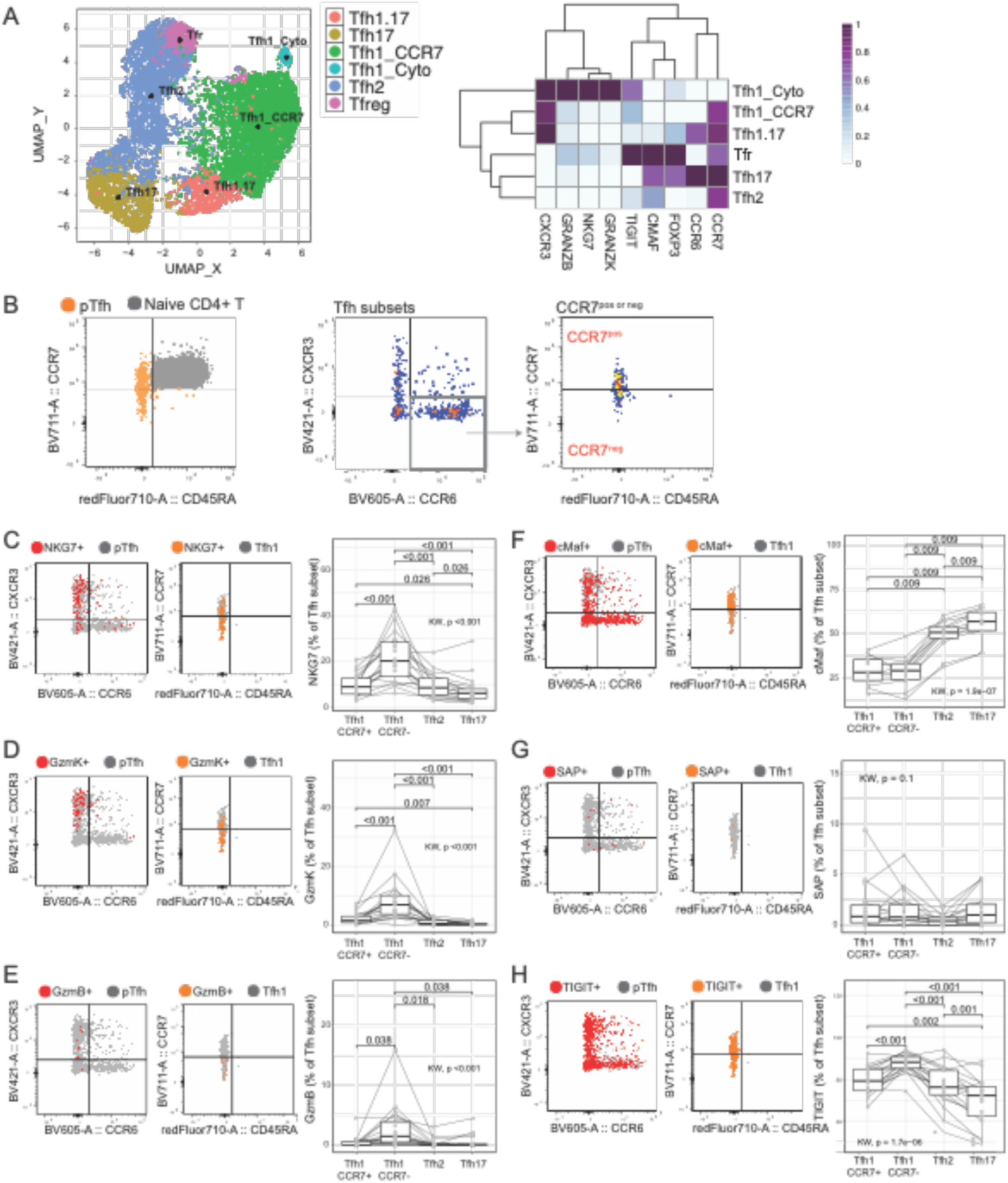
Phenotypic diversity of pTfh cell subsets based on CCR7 expression. A comprehensive spectral flow cytometry panel was used to analyse pTfh cells in healthy individuals (n =19). (**A)** pTfh cells were identified by CXCR5 and PD1 and analysed with unbiased analysis of indicated markers. UMAP of a single experiment analysing n=7 individuals. Analysis performed three independent times. (**B)** Tfh cells were analysed with manual gating. A gradient of CCR7 expression is detected within pTfh cells, and pTfh1 CCR7neg (CCR7-) and pTfh1 CCR7pos (CCR7+) subsets can be identified, along with pTfh2 and pTfh17 cells. Expression of cytotoxic markers NKG7 **(C)**, Granzyme K **(D)** and Granzyme B **(E)** and transcriptional factors cMaf **(F)**, SAP **(G)**, and TIGIT **(H)** were analysed within pTfh cell subsets. Flow plots are a representative individual with marker expression overlayed against CXCR3 and CCR6 or CCR7 and CD45RA. Boxplots with median and IQR are indicated along with individuals. P is the paired Wilcoxon signed-rank test after adjustment with multiple testing correction using Holm’s method, while the Kruskal-Wallis test is used for the global comparison. Only significant differences (p<0.05) are shown. See also Supplementary Fig. 2.

To assess if analysis of a single marker, CCR7, together with CXCR3 and CCR6 could capture this underlying diversity in Tfh1 cells, we next applied manual gating approaches. Within pTfh cells (CXCR5+PD1+ [FoxP3-/CD25-] CD4 T cells) a gradient of CCR7 expression was detected, and pTfh1 cells could be divided into CCR7^pos^ and CCR7^neg^ subsets (**Fig. 2B, Supplementary** Fig. 2D). Within pTfh subsets, expression of cytotoxic markers NKG7, GrzmB and GrzmK was significantly higher in CCR7^neg^ pTfh1 compared to either CCR7^pos^ pTfh1 or pTfh2 and pTfh17 cells (**Fig. 2C-E** **Supplementary** Fig. 2E). Within pTfh cells, we also quantified cMaf which was upregulated in Tfh2 cell clusters, and SAP (*SH2D1A)* and TIGIT which were increased transcriptionally in Tfh1_CCR7 cells (**Fig. 1D**). cMaf and SAP have essential roles in GC Tfh cell development ^34^, and TIGIT is reported to have highest expression in GC-Tfh and pTfh2 cells ^8^. cMaf expression was not different between pTfh1 CCR7^neg/pos^ cells, both of which had significantly lower expression compared pTfh2 and pTfh17, consistent with previous reports ^8^ (**Fig. 2F**). SAP expression was low across all subsets, and was lowest in pTfh2, and comparable between the two pTfh1 subsets (**Fig. 2G**). Further, in contrast to previous reports, TIGIT expression was highest on pTfh1 CCR7^neg^ cells compared to both pTfh1 CCR7^pos^ and pTfh2 cells (**Fig. 2H**). Thus, while CCR7 expression could differentiate populations of pTfh1 cells with distinct cytotoxic potential, how key transcriptional factors may influence pTfh subsets requires further investigation.

To further investigate potential functional differences of pTfh1 cell subsets, cytokine production from pTfh cells was quantified following PMA/Io stimulation in healthy donors (**Supplementary** Fig. 3**, Supplementary Table S4,** n=11, sex 36.36 % males, age 37.5[32-40], median[IQR] years). As seen *ex vivo*, a gradient of CCR7 expression is detected within pTfh1 cell subsets identified with CXCR3 and CCR6 expression post stimulation (**Supplementary** Fig. 3A**/B**). Following PMA/Io stimulation and within the commonly identified pTfh1, pTfh2 and pTfh17 cell subsets, the production of the Tfh-related cytokine IL-21 was comparable, while pTfh1 produced higher IFNγ and TNF, pTfh2 produced higher IL-4 and pTfh17 produced higher IL-17, as expected (**Supplementary** Fig. 3C**/D**). Production of the regulatory cytokine IL-10 was detected at low levels in all subsets. When pTfh1 cells were analysed based on CCR7 expression, there was no difference between pTfh1 CCR7^pos^ and CCR7neg cells in IL-21, or TNF production, however Tfh1 CCR7neg cells produced significantly higher IFNγ, IL-4 and a trend towards increased IL-10 (**Fig. 3A**). Further, when the total cytokine composition was analysed, there was a significant difference in the cytokine composition produced by pTfh1 CCR7^pos^ compared to CCR7^neg^ cells (**Fig. 3B).** Given the importance of the milieu Tfh cytokine production in B cell responses ^5^, data suggest that subsets may have nuanced roles in antibody induction during infection.

**Figure 3.**
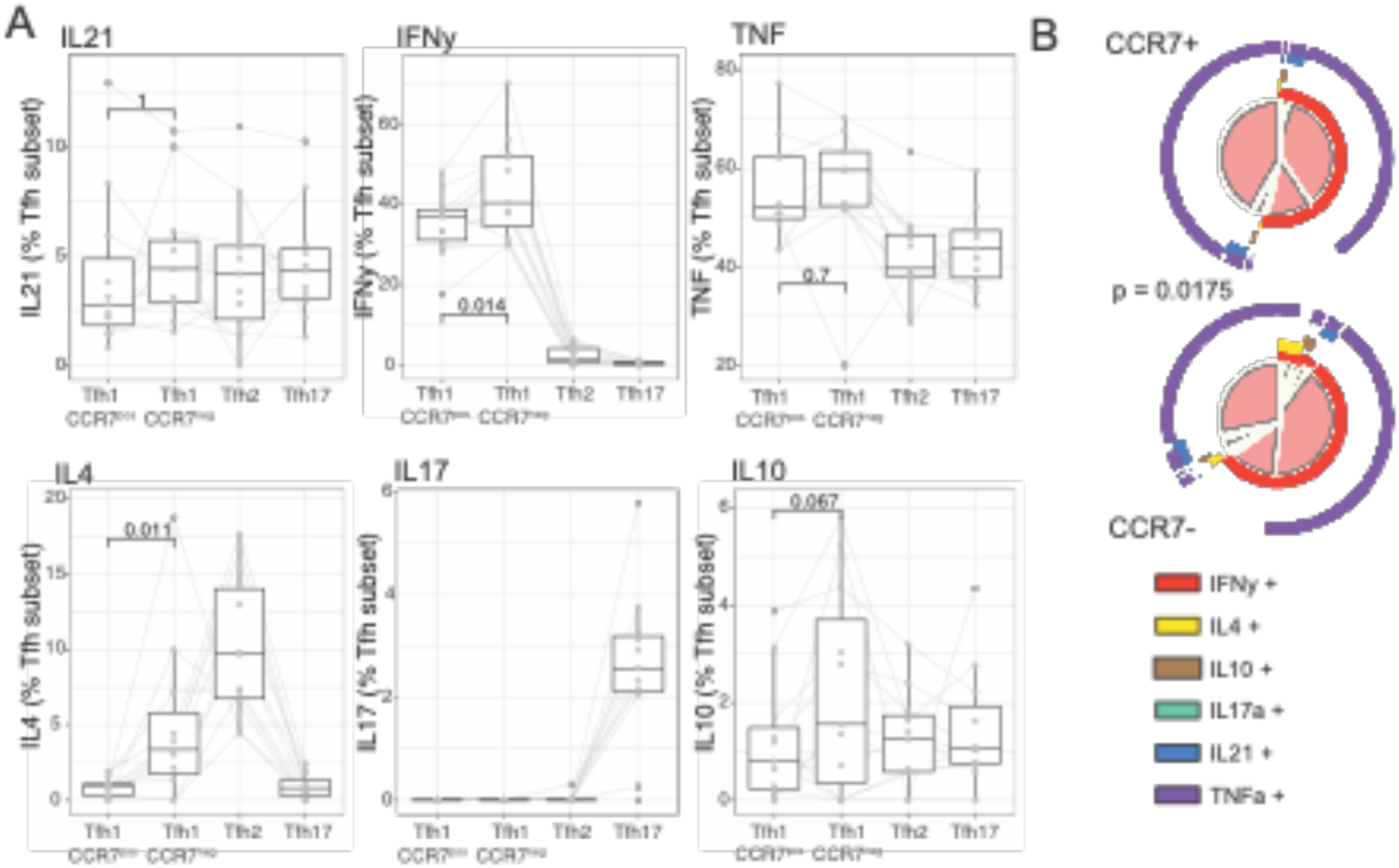
Functional diversity within pTfh cell subsets based on CCR7 expression. Cytokine production following PMA/ Io stimulation was measured by intracellular staining in healthy individuals (n=11). pTfh cells subsets were identified by CXCR3 and CCR6 expression as pTfh1 (CXCR3+CCR6+), pTfh2 (CXCR3-CCR6-) and pTfh17 (CXCR3-CCR6+), and pTfh1 cells subsets identified as CCR7^pos^ and CCR7^neg^ cells. **A)**. IL-21, IFNγ, TNF, IL-4, IL-17, and IL-10 expression within pTfh cell subsets. P is the paired Wilcoxon signed-rank test between pTfh1 cell subsets. **B**) Overall cytokine production within pTfh1 CCR7^pos^ and CCR7^neg^ cells. P is permutation test in SPICE. Box and whisker plots indicate first and third quartiles for hinges, median line and lowest and highest values no further than 1.5 IQR from hinges. All individual data are represented by the grey dots and lines. See also Supplementary Fig. 3.

Together, these data show that transcriptionally, phenotypically and functionally, pTfh1 cell subsets identified by CXCR3+ expression can be further distinguished based on CCR7 expression. These data highlight a previously unrecognised diversity of pTfh cells in healthy individuals when only considering subsets identified by CXCR3 and CCR6 expression, which may have important implication to B cell development during infection.

### Expansion of pTfh1 and pTfh2 cell subsets in human malaria infection

To investigate the roles of pTfh1 CCR7^pos^ and pTfh1 CCR7^neg^ subsets, along with other pTfh cells, during infection, we applied findings to human malaria with samples from a CHMI study cohort. pTfh cells (CD45RA-CXCR5+PD1+) were sorted from individuals in CHMI at day 0 (prior to inoculation), day 8 (peak infection), day 15 and day 36 (both after anti-parasitic drug treatment), and their transcriptional profiles analysed using 5’ droplet scRNAseq (n =4, 100 % males, age 22 [21.5-26], median [IQR] years, **Supplementary** Fig. 4A**/B**). After QC, 27,552 cells were analysed, with high quality cells captured from each donor and timepoint (day 0 n = 6,741, day 8 n=6,125, day 16 n=8,103, day 36 n=7,583, **Supplementary** Fig. 4C). Data were integrated based on donor and time point **(Supplementary** Fig. 4D), and pTfh cell clusters identified by label transfer with scType (an automated cell-type identification tool ^35^), using the top 20 upregulated and downregulated marker genes for each cell cluster identified in the healthy ‘map’ data set used as a reference (**Supplementary Table S5**, **Supplementary** Fig. 4E-H). scType specificity scores were used to collapse clusters into cell subsets (**Figure 4A, Supplementary** Fig. 4H). This approach identified Tfh1_cytotoxic, Th1_CCR7, Tfh2, Tfh17, IFNI, stress, ribosomal, cluster 2 and cluster 3 subsets. Tfh1.17 and Tfr cells were not included in the reference map because of overlapping signatures with Tfh1/Tfh17 and Tfh2 subsets. Following label transfer, Tfh1.17 could not be identified manually, however Tfr cells were identified by inspection of marker genes *FOXP3, CTLA4* and *IL2RA* (**Fig. 4B**). Gene expression of annotated cell clusters was consistent with the reference data set (**Fig. 4B, Supplementary** Fig. 4G**, Supplementary Table S6**). For example, Tfh1_cytotoxic and Tfh1_CCR7 clusters both expressed *CXCR3*, with the two clusters having differing levels of inflammatory markers including *GZMK*, *CCL5* and *NKG7,* which were increased in Tfh1_cytotoxic, and the memory markers *SELL* and *CCR7,* which were increased in Tfh1_CCR7 (**Fig. 4B)**. Similarly, Tfh17 cells identified with label transfer had increased expression of *CCR6*, *BHLHE40* and *TNFRSF4*.

**Figure 4:**
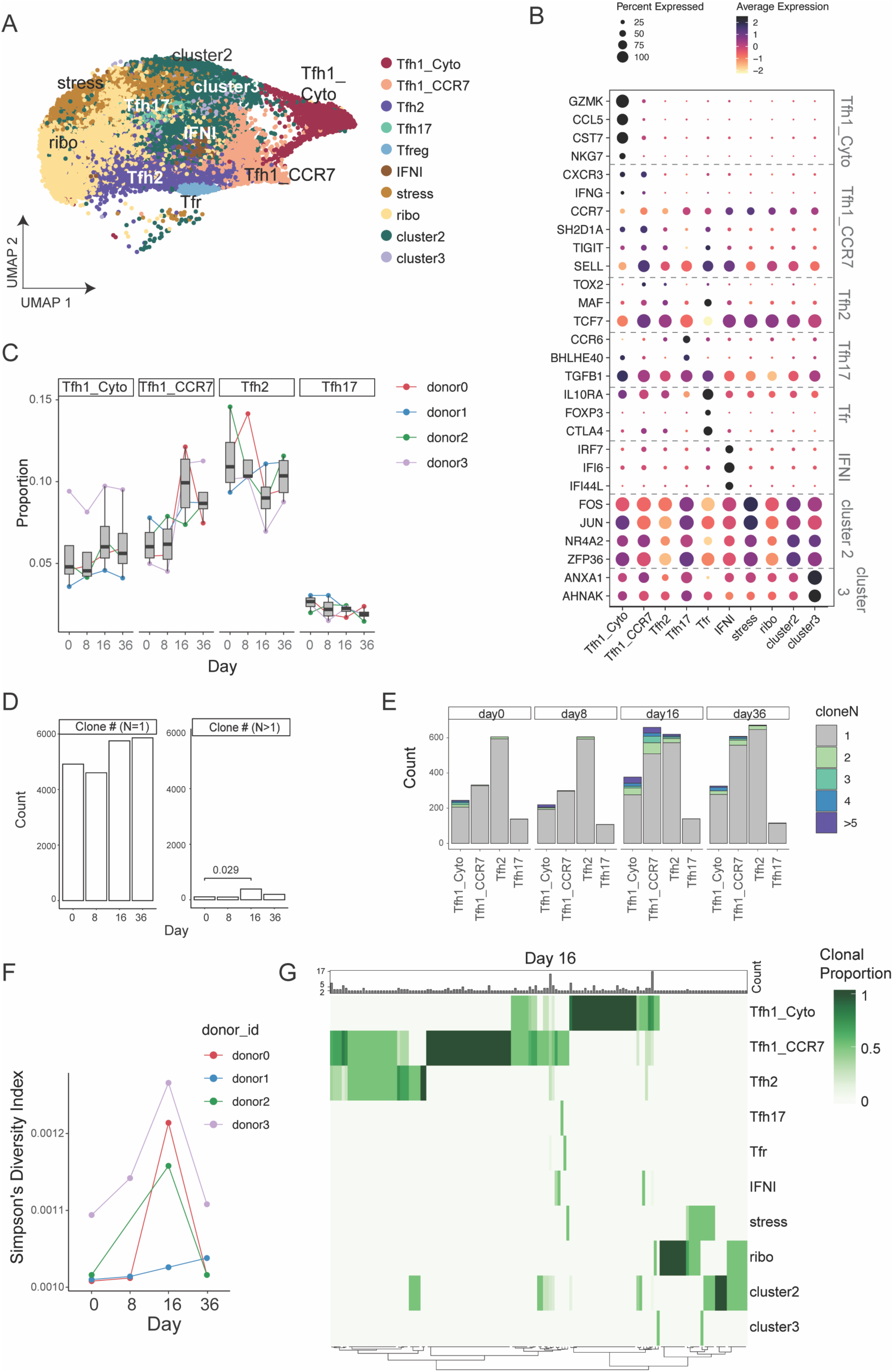
Clonal expansion of diverse pTfh cell subsets during malaria infection. **A)** pTfh cells (CXCR5+PD1+nnCD4 T cells) were isolated from individuals (n=4) during controlled human malaria infection (CHMI) at day 0, 8, 16 and 36. scType label transfer was used to identify Tfh transcriptional subsets using the healthy Tfh cell data set as a reference. **B.** Average expression of marker genes used to annotate reference data set in each subset in CHMI data set **C.** Relative proportion of each pTfh transcriptional cluster during CHMI. Each individual is show. Statistical testing was not performed due to limited sample size in scRNAseq data set. **D.** Counts of clones with either a clone size = 1 (top) or clone size ý 2 across days (bottom). Clonal expansion was detected at day 16. **E.** Clone family size count over time in each subset - Clonal expansion was detected in Tfh1_cyto, Tfh1_CCR7 and Tfh2 clusters at day 16. **F.** Simpsons diversity Index in each donor across time. **G.** Clonal overlap at day 16 across clusters. Clonal size is indicated in count in top bar, each clonal proportion across subsets is indicated. See also Supplementary Fig. 4 and 5.

Interestingly, *TCF7*, *MAF* and *TOX2*, which were identified as marker genes in the reference data set for Tfh2, were expressed both within the Tfh2 subset, but also relatively increased in the Tfh1_CCR7 cells in the infection data set (**Fig. 4B, Supplementary** Fig. 4G). Together, these data show the utility of the reference map data set to identify Tfh cell subsets within sorted pTfh cells during multiple time points in infection.

To investigate how the Tfh cell compartment was impacted by malaria, we assessed the proportion of each cluster over time. There was a clear expansion of the Tfh1_CCR7 subset in all individuals at day 16 post infection, while Tfh1_cytotoxic cells were expanded at this time point to a lesser magnitude (**Fig. 4C**). Tfh2 cells expanded in three of four individuals at day 8, consistent with the earlier activation of this subset in this cohort reported previously ^12^. Other subsets were not clearly impacted by infection (**Supplementary** Fig. 4I). Within each subset, TCRA/B sequences were analysed. TCR sequences with complete TRA/TRB genes were captured for 21,939 (77.97%) cells, and this was comparable across the days/cell types (**Supplementary** Fig. 5A). Clones were defined as cells sharing identical amino acid sequence in both TRA and TRB. During CHMI there was evidence of clonal expansion at day 16, with a significant increase in clonal family size greater than 1 (**Fig. 4D/E**), and increased Simpson Index, indicating increased clonal expansion (**Fig. 4F**). Expansion was highest in Tfh1_cytotoxic and Tfh1_CCR7 clusters, consistent with the increased frequency of these cells and indicative of clonal proliferation (**Fig. 4E, Supplementary** Fig. 5B). Clonal expansion was also detected within Tfh2 cells at day 16 (**Fig. 4E**). As expected, minimal clonal sharing was detected between donors with only a single clone shared between donor 0 and donor 1 on day 36, which may indicate sequence convergence detected due to analysis of clones with amino acid rather than nucleotide sequence ^36,37^ (**Supplementary** Fig. 5C). All other clonal sharing occurred within an individual, the majority at a single timepoint (**Supplementary** Fig. 5D). For donor 3, clones were also shared across time points including day 0 prior to malaria, with large clonal families maintaining a single fate across time in the Tfh1_cytotoxic cell cluster, possibly indicating a subset expansion during a prior infection (**Supplementary** Fig. 5D). At day 16, where malaria associated expansion was most evident, clonal sharing across Tfh cell clusters were analysed. While there was a minimal sharing between Tfh1_cytotoxic and Tfh2 cell subsets, Tfh1_CCR7 cells shared clones with both Tfh1_cytotoxic and Tfh2 cells, consistent with the overlapping profiles of Tfh1_CCR7 with both Tfh1_cytotoxic and Tfh2 subsets (**Fig. 4G**). Taken together, these data are consistent with the emergence of malaria specific Tfh cells with diverse transcriptional signatures during infection. These include pTfh1 CCR7^pos^ cells with increased functional capacity, pTfh1 CCR7^neg^ cells with cytotoxic phenotypes, and pTfh2 cells which are transcriptionally most similar to GC Tfh cells ^8^.

### Unique transcriptional activation of Tfh1 subsets during human malaria infection

To investigate transcriptional pathways activated in each Tfh transcriptional subset during malaria, differential gene expression between day 0 and subsequent time points was performed with pseudo- bulk analysis, grouped by individual (**Supplementary Table S7**). The largest transcriptional changes were detected at day 16 within the Tfh1_CCR7 and Tfh1_cytotoxic subsets (**Fig. 5A**). Differentially expressed genes (DEGs) at day 16 were also detected in Tfh2 cells consistent with clonal expansion of these cells, and previously reported findings ^12^ (**Fig. 5A**). The number of DEGs was contracted by day 36, with DEGs detected at this time point restricted to Tfh1_CCR7, suggesting this subset had a sustained response to infection. The majority of DEGs at day 16 from Tfh1_CCR7, Tfh1_cytotoxic, and Tfh2 were subset specific (**Fig. 5B**). Shared across all three subsets was the activation marker *CD38*, and Tfh1_cytotoxic and Tfh1_CCR7 cells also both upregulated *MKI67* consistent with proliferation and clonal expansion of these cells during malaria. However, overall expression of activation and proliferation genes was highest in Tfh1_CCR7 cells (**Fig. 5C, Supplementary** Fig. 6A). Analysis of the top 20 upregulated genes from Tfh1_CCR7, Tfh1_cytotoxic and Tfh2 cells highlighted the uniquely upregulated gene signatures identified from Tfh1_CCR7 and Tfh1_cytotoxic cells (**Fig. 5D, Supplementary** Fig. 6D). Additionally, while shared upregulated genes also included genes associated with increased inflammation and cytotoxicity including *NKG7, CCL5, GZMH, CXCR3* and *IFNG* the level of these genes was significantly higher in Tfh1_cytotoxic cells (**Fig. 5D**, **Supplementary** Fig. 6B). Further, the transcriptional profile of Tfh2 cells was also maintained and similar to that seen at day 0, despite upregulation of Tfh1 associated genes (**Supplementary** Fig. 6C).

**Figure 5:**
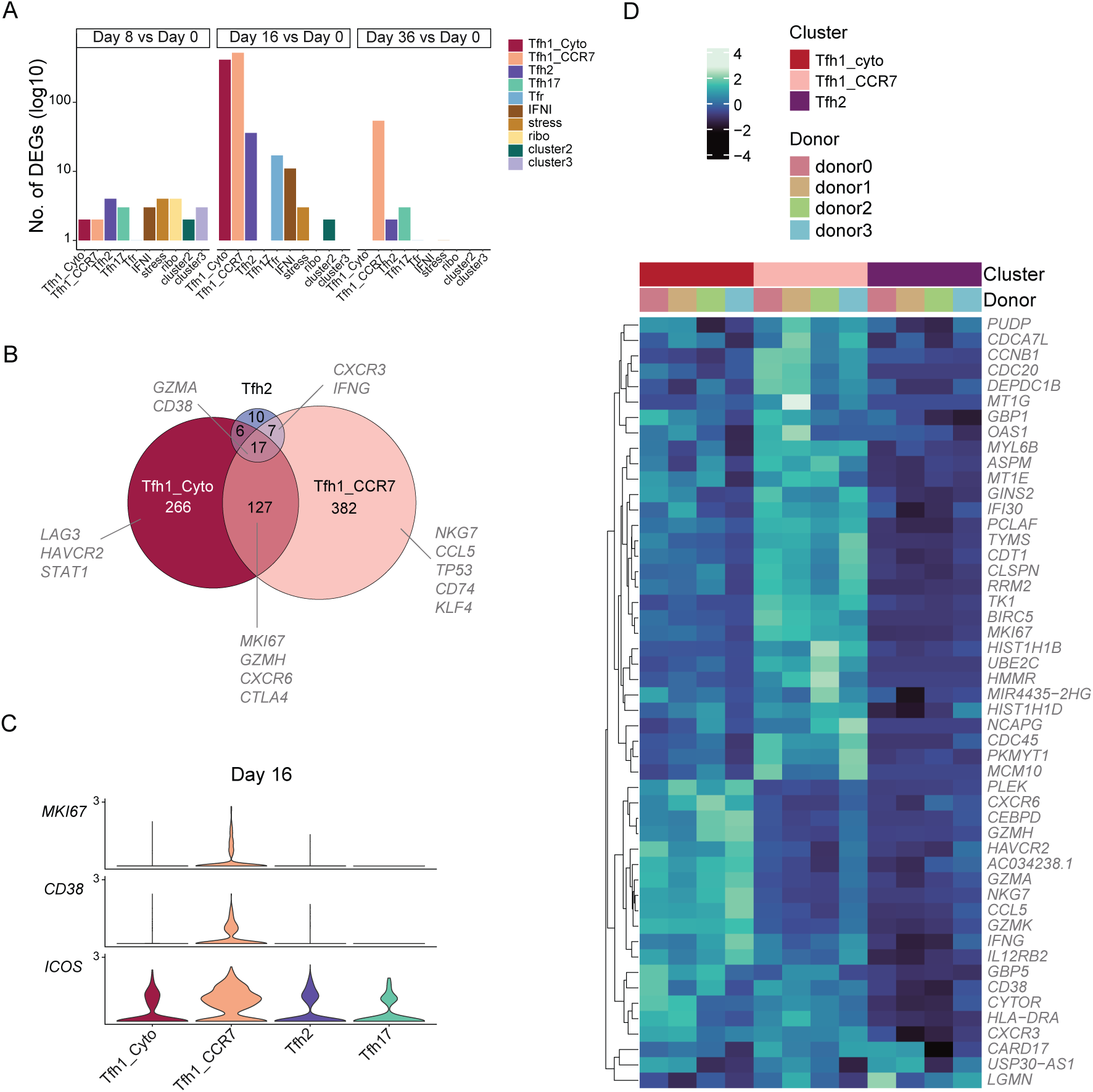
Transcriptional activation of Tfh cell subsets during controlled human malaria infection. **>A.** DEGs for each cluster were calculated at day 8, 16, and 36 compared to day 0. The number of DEGs for each subset and timepoint. **B.** Venn diagram indicating the shared and unique DEGS at day 16 for Tfh1_cytotoxic, Tfh1_CCR7 and Tfh2 subsets. **C.** Average expression of activation genes MKI67, CD38 and ICOS at day 16 in each Tfh subset compared to Day 0. **D.** Average expression of top 50 unique upregulated genes for Tfh1_Cyto, Tfh1_CCR7 and Tfh2 (curated from top 20 upregulated DEGs in Tfh1_CCR7, Tfh1_cytotoxic and Tfh2) cells across all three subsets at day 16 subsets. See also Supplementary Figure 6.

Taken together, scRNAseq analysis of pTfh cells during malaria identify the activation, proliferation and clonal expansion of Tfh1_CCR7, Tfh1_cytotoxic and Tfh2 cells with unique transcriptional activation profiles.

### Frequency and proliferation of malaria specific pTfh CCR7^pos^ cells is associated with the magnitude and function of antibody induction in CHMI

To confirm transcriptional findings at the phenotypic and antigen specific level and assess associations of different pTfh subsets with antibody induction, we applied our flow cytometry panel to identify pTfh subsets, including pTfh1 CCR7^pos^ and CCR7^neg^ cells to assessed the activation of pTfh cells during CHMI in a subset of individuals from a previously published cohort (n= 19, sex 83.3%% males, aged 24 years [20.2-28.2, median [IQR] years) ^12^. Here we expanded on the published analysis by focusing on malaria specific pTfh cells which were identified by Activated Induced Marker (AIM) assays following PBMC stimulation with parasite infected RBCs (pRBCs) ^16^ (**Supplementary** Fig. 7A). Consistent with scRNAseq analysis, AIM+ (CD69+OX40+) pTfh cells were expanded at day 16 and 36 during CHMI. The frequencies of cells detected in control stimulated cultures (uninfected RBCs) were unchanged (**Fig. 6A).** The largest response of malaria specific (AIM+ in pRBC stim) pTfh cells occurred in pTfh1 CCR7^pos^ cells, however, increased frequencies of Tfh1 CCR7^neg^ and Tfh2 cells were also detected, consistent with the proliferation and expansion of these cells detected transcriptionally (**Fig. 6B**). Proliferation of AIM+ pTfh17 cells were also detected at day 36 (p=0.051), however given that pTfh17 cells did not expand clonally, nor have any detectable transcriptional changes during infection, these cells are likely to be TCR independent cells which are activated via cytokines within AIM assays ^38^. Within malaria specific pTfh1 CCR7^pos^, pTfh1 CCR7^neg^ and pTfh2 cells, there were significant increases in expression of activation markers ICOS and CD38, and proliferation marker Ki67 at day 16 and 36 (**Fig. 6C/D**). However, the expression of these markers was significantly higher on malaria specific pTfh1 CCR7^pos^ cells compared to other subsets, consistent with the greatest expansion and activation of pTfh1 CCR7^pos^ cells transcriptionally (**Fig. 6E/F**).

**Figure 6:**
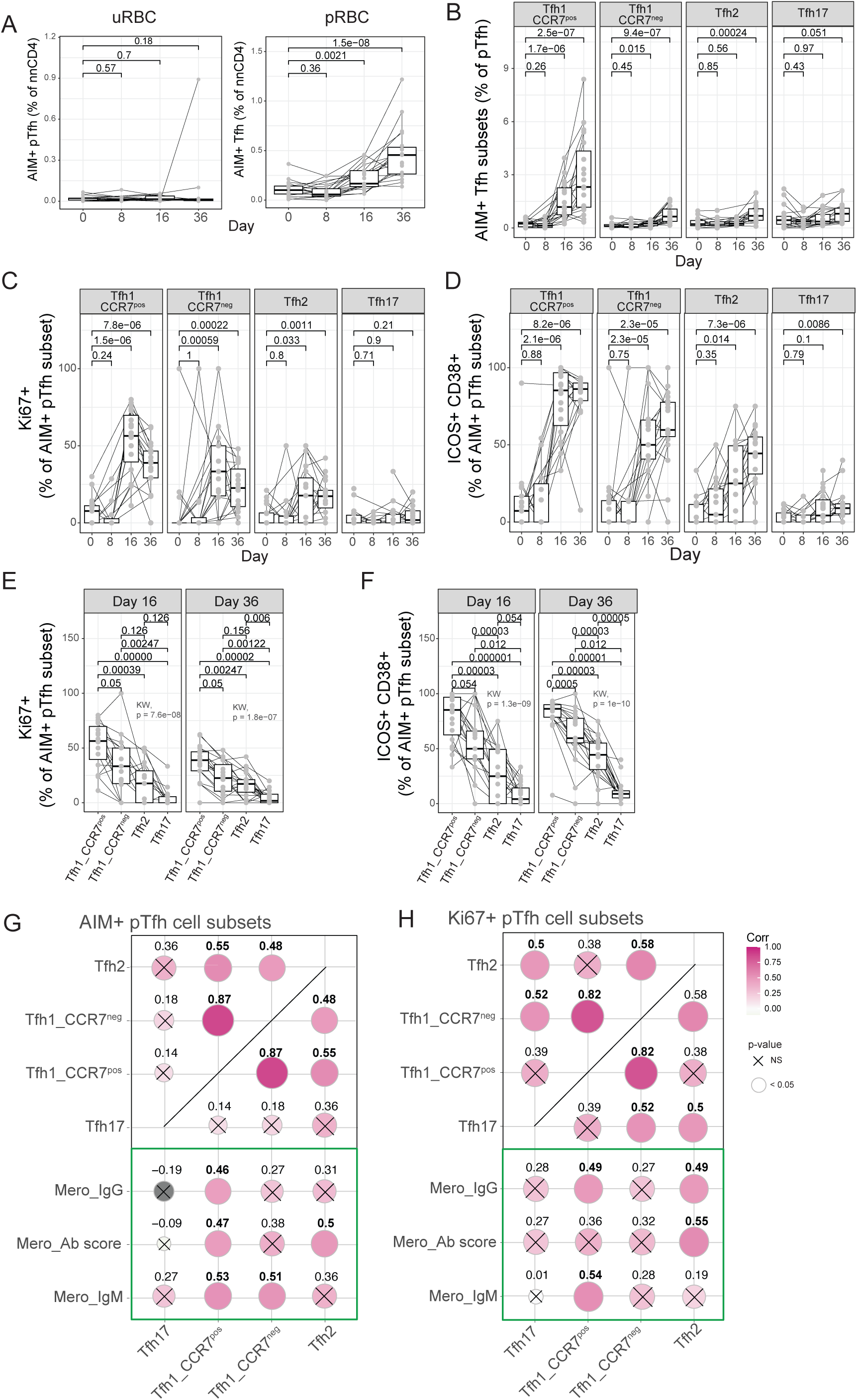
Activation of malaria specific Tfh cells during malaria infection and association with antibody induction. **A.** Activation induced marker (AIM) assays were used to detected malaria specific pTfh cells in individuals following CHMI (n= 15-20). AIM+ cells were identified as CD69+OX40+ pTfh cells after stimulation with uninfected or parasite infected RBCs (uRBC and pRBC respectively) within all non- naïve CD4 T cells. Malaria specific Tfh cells increased at day 16 and day 36 after infection. **B.** Malaria specific (AIM+) subset specific pTfh cells as a proportion of total pTfh cells. **C-D.** Ki67+ (C) and ICOS+CD38+ (D) cells as a percentage of each AIM+ malaria specific pTfh subset during CHMI comparing changes with infection. **E-F.** Ki67+ (E) and ICOS+CD38+ (F) cells as a percentage of each AIM+ malaria specific pTfh subset at day 16, comparing expression levels between subsets. **G-H.** Correlation between (**G)** malaria specific pTfh cell subsets or (**H)** Ki67 expression of Tfh subsets at day 36 with total IgG and IgM levels to merozoite, and overall antibody score against merozoite, which captures the total antibody magnitude, breadth and functionality specific to merozoite. **A-D**. P is the Wilcoxon rank-sum test for each infection timepoint against baseline. **E-F**. P is the Wilcoxon rank-sum test after adjustment with multiple testing correction using Holm’s method, while the Kruskal-Wallis test is used for the global comparison. **G-H**. Spearman’s rho and p-value are indicated. See also Supplementary Figure 6 and Supplementary Figure 7.

To investigate the link between each malaria specific pTfh cells and antibody development, we correlated malaria-specific pTfh cells with antibodies produced after infection in response to intact merozoites, quantified previously ^12^. Antibody development was quantified with an antibody score which captured the magnitude, breadth and functionality of response, by measuring IgM, IgG and IgG subclasses, C1q fixation, FcR crosslinking and opsonic phagocytosis ^12^. The frequencies of malaria- specific pTfh1 CCR7^pos^, pTfh1 CCR7^neg^ and pTfh2 cells correlated with each other, but not with pTfh17 cells (**Fig. 6G)**. Despite this, pTfh1 CCR7^pos^ were correlated the most strongly and consistently with antibody induction compared to other subsets, and were positively correlated with total antibody score, and IgM and IgG to the merozoite (**Fig. 6G**), along with IgG1 and IgG2 (**Supplementary Fig. S7B).** In contrast, pTfh1 CCR7^neg^ cells only correlated with IgM and pTfh2 cells only correlated with antibody score (**Fig. 6G**). Similarly, while Ki67 expression amongst subsets was correlated, only Ki67+ pTfh1 CCR7^pos^ and Ki67+ pTfh2 cells were correlated with antibody score, IgG and IgM and antibody functions (**Fig. 6H, Supplementary Fig. S7D/E)**. Together, data highlight that pTfh1 CCR7^pos^ and pTfh1 CCR7^neg^ cells have distinct functional potential. While both are activated in malaria, pTfh1 CCR7^pos^ cells, together with pTfh2 cells, are more strongly associated with antibody development.

## Discussion

Here we provide data that advances understanding of Tfh cell subsets, highlighting their functional diversity across the course of human malaria. Using unbiased analysis to integrate pTfh in the resting state, we identify two clearly distinct pTfh1 subsets, which we show can be distinguished based on CCR7 expression phenotypically. pTfh1 CCR7^pos^cells do not express cytotoxic markers and produce less inflammatory cytokines, but similar IL-21 levels compared to pTfh1 CCR7^neg^ cells. Expanding our findings to the context of human infection, we show during malaria that while pTfh1 CCR7^pos^, pTfh1 cytotoxic/CCR7^neg^, and Tfh2 cell subsets expand, and are activated, activation pathways are unique between the subsets. We show both transcriptionally and at the malaria specific level, the magnitude of activation is greatest in pTfh1 CCR7^pos^ cells, which also expand clonally. Additionally, the frequency and proliferation of pTfh1 CCR7^pos^ cells was most consistently associated with antibody induction compared to other subsets. Taken together, these findings provide new knowledge on Tfh cell activation and functions during human infection. The specific role of pTfh1 CCR7^pos^ and pTfh2 cells, but not pTfh1 CCR7^neg^ responses in antibody induction during malaria informs approaches to target Tfh cells to improve antibody development and protection.

Understanding the mechanisms driving the development of distinct Tfh cell subsets has important translational implications for vaccine design and therapeutic interventions. Development of distinct Tfh cell subsets is suggested to be mediated by cytokine skewing driven by distinct inflammatory milieu occurring during different immune responses to infection or vaccination ^5^. A recent investigation of Tfh cell development in mice immunized with Type-1 or Type-2 adjuvanted conditions supported this concept, inducing Tfh1 or Tfh2 cell-dominated responses, respectively ^32^. However, a comparative investigation of Tfh cell development and function in different infections has highlighted that cytokine signalling within Tfh cells is more important in driving subset fate than pathogen class ^7^. For example, within that study Tfh cell fates induced by LCMV and influenza, both of which drive type-1 immune responses, were distinct and drove differing antibody responses ^7^, highlighting nuanced roles of Tfh cell subsets in pathogen specific antibody development. While Tfh cells enriched for different cytokine signalling pathways could be identified in human tonsils ^7^, how distinct Tfh cell subsets develop in human infection is unclear. Further, our data highlight that even within a single infection, induced pTfh cells are diverse. During malaria infection both pTfh1 CCR7^pos^ and CCR7^neg^ subsets, along with pTfh2 are expanded, transcriptionally activated and malaria specific. pTfh1 CCR7^pos^ cells shared clonal overlap with both pTfh1 CCR7^neg^ cells and pTfh2 subsets, but these later subsets were largely clonally distinct from each other. Shared clonal relationships of malaria specific pTfh1 and pTfh2 cells has been reported previously ^39^. Whether these relationships indicate cells along a single developmental pathway, or divergence from a parent population and how cytokine signalling may influence subsets development is unknown.

The importance of understanding Tfh cell subset development is highlighted by the distinct functional consequences on antibody development linked to Tfh cell subset function. Human studies linking Tfh cell subsets to antibody development have largely been limited to identifying Tfh cell subsets based on CXCR3 and CCR6 expression ^8,9,40,41^, including in malaria ^12,16,17,19,39^. Alternatively, studies which include a more comprehensive analysis of pTfh cells have not related Tfh cell diversity to antibody development ^31,42^. Here, our scRNAseq analysis shows that while CXCR3 and CCR6 are informative, diversity is masked by this gating strategy. Of functional relevance, within pTfh1-like cells, we found that pTfh1 CCR7^neg^ cells contain the large majority of Tfh cells expressing cytotoxic markers, including *NKG7,* granzyme K and granzyme B. In contrast, these markers were absent from Tfh1 CCR7^pos^ cells, which produce less inflammatory cytokines but comparable levels of IL-21, resulting in distinct cytokine milieu. Tfh cells with expression of cytotoxic markers have been identified previously, and GC-Tfh cells which produce granzyme B in response to group A *Streptococcus* kill B cells, potentially hampering antibody development ^43^. More recently granzyme K+ CMV specific pTfh cells were reported in humans. However, the function of these cells in the context of antibody development is unknown ^32^. During malaria infection, we show that that while malaria specific pTfh1 CCR7^pos^, pTfh1 CCR7^neg^ and pTfh2 cell subsets are induced, only pTfh1 CCR7^pos^ and pTfh2 cell subsets were clearly associated with antibody development. We have previously reported in the same cohort that global pTfh2 cell activation early in infection (not considering antigen specificity) was associated with antibody development ^12^. While the importance of pTfh2 cells in antibody development to malaria has been supported by others ^13–15^, a negative role of pTfh1 in antibody development was difficult to rationalise in the context of what is known from *in vitro* studies and animal models of induction of cytophilic IgG3 antibodies which protect from malaria ^21^. Further, our data from children in Ugandan in a high malaria transmission setting showed that children who had the highest levels of functional antibodies had increased proliferating pTfh2 and pTfh1 cells ^19^. Data here resolve these contradictions with the identification of distinct pTfh1 cell subsets. Further studies are required to investigate whether pTfh1 CCR7^neg^ cells with cytotoxic functions have negative roles in antibody induction, and how pTfh1 CCR7^pos^ and pTfh2 cells function to induce protective antibodies in malaria.

Limitations of our study include the analysis of pTfh cells isolated from peripheral blood of volunteers, and not GC-Tfh within secondary lymphoid tissues. Future approaches to obtain and analyse human secondary lymphoid tissue in the context of health and disease are needed. While pTfh cells share overlapping signatures with GC-Tfh cells and are thought to provide a snapshot of the GC response ^6^, activated pTfh may not fully represent all GC-Tfh phenotypes and functions ^44^. Future studies will focus on developing approaches for fine needle aspirate studies of draining lymph nodes during human malaria infection as applied to COVID and influenza vaccination settings ^44–48^ , and investigating the immune response within the spleen in malaria infected individuals which is likely a key site of antibody development in blood stage malaria ^49,50^. Nevertheless, such approaches are not feasible for large human cohorts, thus data here on pTfh cells provides important insights that can be applied to other human infection and vaccinations. Future studies dissecting mechanisms of protective and disruptive Tfh cell development may lead to avenues to modulate Tfh cells during infection or vaccination to induced more robust and protective antibody responses.

In conclusion, this study provides important new knowledge on functional diversity with pTfh cells during human infection. We identify specific pTfh cell subsets that associate with antibody induction in malaria, thus informing the development of approaches that can target these cells to improve antibody development induced during vaccination. We provide a scRNAseq data set that can be applied to other diseases, and a optimised approach to identify distinct pTfh1 subsets based on CCR7 expression. The identification of further Tfh subsets and their potential functional importance is highly relevant to developing novel therapeutics or new vaccine strategies that can induce potent and long-lasting immunity against not only malaria, but other important pathogens.

## Materials and methods

### Ethics statement

Ethics approval for the use of human samples was obtained from the QIMR Berghofer Human Research Ethics Committee (HREC P1479) and the Alfred Health Ethics Committee (288/23). Written informed consent was obtained from all study participants.

### Study cohorts

For CHMI studies, inoculum preparation, volunteer recruitment, infection, monitoring and treatment were performed as previously described ^51^. In brief, healthy malaria naive individuals underwent CHMI with 2800 viable *P. falciparum* parasitized RBCs, and peripheral parasitaemia was measured at least daily by qPCR as described previously ^52^. Participants were treated with antimalarial drugs at day 8 of infection when parasitaemia reaches approximately 20,000 parasites/ml. Blood samples (from 5 studies across 6 independent cohorts) were collected prior to infection (day 0), at peak infection (day 8) and 14 or 15 and 27-36 days (end of study, EOS) after inoculation (in analyses these time points are grouped as 0, 8, 14/15 and EOS). PBMCs were isolated by Ficoll-Paque (Sigma, USA) density gradient centrifugation, isolated PBMCs were cryopreserved in 10% DMSO/FBS. Participants were healthy malaria naïve adults with no prior exposure to malaria or residence in malaria-endemic regions. Clinical trials were registered at ClinicalTrials.gov NCT02867059 ^53^, NCT02783833 ^54^, NCT02431637 ^55^, and NCT02431650 ^55^. For trials NCT02867059, NCT02783833, NCT02431637 all participants received the same study drug (no-randomization) and 3D7-strain parasites. For NCT02431650, participants were randomized 1:1 to receive study drug or control and 3D7-strain parasites. Samples used here were from samples opportunistically collected from volunteers who consented to donate blood for immunological studies within the parent clinical trial. As such, no sample size estimation was performed for this immunology study. PBMCs from healthy non-infected controls was collected by the same processes for analysis of responses following PMA/Io stimulation, development of transcriptional map of pTfh cells in health donors.

*Sex as a biological variable:* Both male and females were included in this study. For some parent CHMI studies females of child baring age were excluded and as such the sex distribution is not evenly distributed.

### Ex vivo cell phenotyping with flow cytometry and activation induced marker assays

PBMCs were thawed in RPMI 1640 (Gibco) containing 10% FCS and 0.02% Benzonase and rested for 2 hours at 37°C, 5% CO2, and then cultured overnight in 96-well plates in 10% FCS/RPMI at 1 x 10^6^ cells per well and stimulated for 18 hours with pRBCS or uninfected red blood cells (uRBCs) at 1:1 ratio. pRBC stimulated cells were used to assess malaria specific responses, and uRBC stimulated cells were used to phenotype all Tfh cells. PBMCs were stained with antibodies against CD49b and LAG-3 along with Fc block for 45 minutes at 37C, 5% CO_2_. Following 2 washes with PBS, cells were then stained with LIVE/DEAD Fixable Blue Dead Cell Stain for 15 minutes at room temperature, washed twice with 2% FCS/PBS, and then surface stained for 15 minutes at room temperature with fluorescent-tagged antibodies from Biolegend (CA, USA), and BD Biosciences (CA, USA) (**Supplementary Table S3**) e. After 2 washes with 2% FCS/PBS Cells were fixed and permeabilized with eBioscienceTM Foxp3 / Transcription Factor Staining Buffer Set (Thermo Fisher Scientific (MA, USA); Cat. #00-5523-00) solution for 20 minutes on ice. After 2 washes with 1X perm buffer, intracellular staining was performed for 30 minutes on ice with fluorescent-tagged antibodies from Biolegend (CA, USA), and BD Biosciences (CA, USA) (**Supplementary Table S3**). Cells were re- suspended in 2% FCS/PBS, and events were collected on the Cytek Aurora 5 laser cytometer (CA, USA).

### PMA stimulation and intracellular cytokine staining

Approximately 1 x 10^6^ PBMCs were used per patient per condition/ stain. Thawing of cryopreserved PBMCs was done as previously described. For cytokine induction following PMA and Ionomycin stimulation, PBMC samples were rested overnight in 10% FCS/RPMI after thawing. After rest, PBMCs were stimulated with PMA (25 ng/ mL; Sigma) and Ionomycin (1 ug/mL; Sigma) for 6 hours at 37°C, 5% CO2. Brefeldin A (10 ug/mL; BD Biosciences) and monensin (10 ug/mL; BD Biosciences) were added after the first 2 hours of stimulation. To account for downregulation of chemokine receptors after exposure to BFA and monensin, antibodies against CCR7, CXCR3, CXCR5 and CCR6 were added alongside Fc block during stimulation with PMA/Ionomycin, prior to addition of BFA and monensin. Anti-CD107a was also included during stimulation for intracellular capture. After stimulation, cells were washed with PBS and stained at room temperature for 15 minutes with LIVE/DEAD Fixable Blue Dead Cell Stain, washed twice with 2% FCS/PBS, and then surface stained with the antibodies listed in **Supplementary Table S4** for 15 minutes at room temperature. Following 2 washes with 2% FCS/PBS, cells underwent fixation/permeabilization with eBioscience Fixation/Permeabilization solution for 20 minutes on ice and washed in permeabilization buffer. Intracellular staining (ICS) with the antibodies listed in **Supplementary Table S4** was performed for 30 minutes on ice following washes with permeabilization buffer and then fixed with BD stabilizing fixative. Samples were acquired on Cytek Aurora 5.

### Flow cytometry data analysis

Flow cytometry data was analysed using FlowJo v10 and then analysed on R (version 4.4.2). For unsupervised data analysis, spectre (version 1.2.0) was used ^56^.

### Single Cell RNA sequencing and analysis

#### Cell isolation

<u>Healthy reference data set –</u> PBMCs from four healthy donors were thawed in thawed in RPMI 1640 (Gibco) containing 10% FCS and 0.02% Benzonase. Cells were rested for 2 hours in 10% FCS/PBS at 37C, 5% CO_2_. After rest, untouched CD4+ T cells were enriched from PBMCs using the CD4+ T cell isolation kit (Miltenyi). Enriched CD4+ T cells were then incubated with Fc block for 15 minutes and then surface stained with antibodies listed in **Supplementary Table S8** for 15 minutes at room temperature. Cells were washed with 0.5% BSA/PBS and stained for viability with Sytox Blue (Invitrogen (CA, USA), Cat. # S34857) and sorted on BD FACSAria™III Cell Sorter into 0.5% BSA/PBS. Four populations of cells were sorted, PD1- Tfh (nnCD4/CXCR5+/PD1-), pTfh1 (nnCD4/CXCR5+/PD1+/CXCR3+ CCR6-), pTfh2 (nnCD4/CXCR5+/PD1+/CXCR3- CCR6-) and pTfh17 (nnCD4/CXCR5+/PD1+/CXCR3- CCR6+). After sorting, cells of a single phenotype were pooled and run on a single 10X Chromium lane.

<u>CHMI –</u> PBMCs collected at day 0, 8, 16, 36 from four participants in CHMI study NCT02867059 ^53^ were thawed in RPMI 1640 (Gibco) containing 10% FCS and 0.02% Benzonase. Cells were washed with 0.5% BSA/PBS and then incubated with Fc block and antibodies against pan-gD and CXCR5 for 15 minutes at room temperature. Following that, the other antibodies listed in Stain 2 in **Supplementary Table S9** were added and cells were incubated for another 15 minutes at room temperature. Cells were washed and stained for viability with Sytox Blue (Invitrogen (CA, USA), Cat. # S34857) and sorted on BD FACSAria™III Cell Sorter into 0.5% BSA/PBS. Live Tfh cells were sorted as nnCD4, CXCR5+PD1+ cells. After sorting, cells from a single time point were pooled and run on a single 10X Chromium lane.

### 10 X Genomics Chromium GEX Library preparation and sequencing

Sorted cells were loaded into each lane of Chromium Next GEM Single Cell 5’ Reagent Kit v2 and Gel Bead-in- Emulsion (GEMs) generated in the 10X Chromium Controller (Chromium Next GEM Single Cell 5’ Kit v2, PN-1000263, Chromium Single cell V(D)J Amplification Kits: Human TCR, PN-1000252, Chromium Next GEM Chip K Single Cell Kit, PN-1000286, , Dual Index Kit TT Set A, PN-1000215). 5’ Gene Expression Libraries and VDJ libraries (TCR) were generated according to the manufacturer’s instructions. Generated libraries were sequenced in a NextSeq 550 System using High Output Kit (150 Cycles) version 1 according to the manufacturer’s protocol using paired-end sequencing (150-bp Read 1 and 150 bp Read 2) with the following parameters Read 1: 26 cycles, i7 index: 10 cycles, i5 index: 10 cycles and Read 2: 90 cycles.

#### Demultiplexing and analysis

ScRNAseq data was demultiplexed, aligned and quantified using Cell Ranger version 3.1.0 software (10x Genomics) against the human reference genome (GRCh38-3.0.0) with default parameters. Cell genotype was obtained using cellsnp-lite ^57^ with the list of informative SNPs obtained from 1000_Genome_Project ^58^. Then vireo was used to predict the original donor utilising the cell’s haplotype ratios, only predicted singlets were used for the analysis ^59^. Cell ranger count matrices for each sample were loaded, merged and analysed using Seurat package v4.3.0 and v5.1.0 ^60^ and visualised with scCustomize v3.0.1 ^61^. The healthy reference and CHMI datasets were processed as two separate datasets. Cells were filtered to remove those expressing fewer than 200 genes and more than 4000 genes, and those with more than 10% mitochondrial content. Only genes expressed in three or more cells were considered. TRA/TRB gene expression were removed from the GEX dataset prior to further downstream processing.

For <u>Healthy Reference map</u> – Dataset was split by donor and then each individual donor dataset was normalised with SCTransform where highly variable genes were also selected. Before integration, the most variable genes shared among the donor datasets were identified using FindintegrationAnchors and were used to integrate the datasets using the IntegrateData function. Following that, PCA dimensionality reduction was performed using RunPCA and the top 20 PCs were used to compute UMAP visualisation using RunUMAP.

For <u>CHMI data</u> - Dataset was split per donor and day, then each individual donor_day dataset was normalised with SCTransform where highly variable genes were also selected.^62^ Before integration, the most variable genes shared among the datasets were identified using FindIntegrationAnchors and used to integrate the datasets using the IntegrateData function. Following that, PCA dimensionality reduction was performed using RunPCA and the top 20 PCs were used to compute UMAP visualisation using RunUMAP.

#### Cell cluster annotation

<u>Healthy reference map –</u> We applied unsupervised clustering using the Louvain algorithm as our modularity optimisation technique and set the granularity of clusters at a resolution of 0.3. A cluster with high mitochondrial content and low read counts was removed as it implies a cluster of cells of low quality. RNA-level counts were log-normalised using NormalizeData (scale factor = 10,000 by default) and were used to run FindMarkers for identifying cluster marker genes as part of cluster annotation. This identified the Tfh1 cluster, along with Tfh2, Tfh17 stress, ribosomal, Type I IFN, cluster 2 and cluster 3 clusters. We then visualised the expression of Treg-related genes (*FOXP3* and *IL2RA)* on the map to identify a cluster of cells belonging to Tfreg within Tfh2 cluster, and another subset of Tfh1/17 cluster which expressed *RORC* along with *CXCR3*, that were positioned separately from the main Tfh1 cluster. Further sub-clustering was also attempted on each of the main Tfh1/2/17 clusters separately to identify any higher-level annotations/groupings that could exist within each subset, however only 2 sub-clusters within the Tfh1 subset were suggested to be of functional difference. Automated cell labelling tool, scType ^35^, was used to transfer labels from the healthy reference Tfh map onto a published human tonsil Tfh dataset ^26^ containing non-GC and GC Tfh subsets. Detailed steps of how the label transfer was performed is as per described below in CHMI data section. Cluster markers of each cluster were further analysed for pathway enrichment using the Ingenuity Pathway Analysis (IPA) suite (v01-22-01, QIAGEN, DUS, Germany).

<u>CHMI data –</u> Unsupervised clustering was calculated using Louvain algorithm with resolution set at 2.2 to achieve overclustering. Low-quality clusters with high mitochondrial content and low read counts were removed. Label transfer using cell type annotations from the Healthy Reference Map onto the CMHI dataset was performed using scType ^35^. The top 20 up- and down-regulated cluster marker genes from each reference cell type were used to calculate a specificity score to predict the cell type for each cluster in the CHMI dataset. Based on the final top-specificity score, similar clusters are then stitched together for the final annotation. As an additional quality check step, a fold-change between the score of the top-predicted cell type versus every other possible cell type was calculated per cluster, and any cluster where the difference between the top-predicted cell type and the next possible cell type was less than 1.1 was flagged as “low confidence”. For clusters 14, 19 and 22 where the top-predicted cell type was flagged as “low-confidence”, a manual comparison of key cell type marker genes was done to confirm the final cell annotation used (final annotation: cluster 14 as Tfh1_CCR7, cluster 19 as cluster 2, cluster 22 as Tfh1_Cyto). After the final prediction, we visualised Treg markers (*“Il2ra”, “Foxp3”*) and manually selected for an area on the UMAP with high expression of these markers as “Tfr” cluster. For the reference database, we omitted Tfh1/17 and Tfr as input as these 2 clusters had signatures commonly shared with the other clusters: Tfh1/17 with Tfh1 and Tfh17, and Tfreg with Tfh2. RNA-level counts were log-normalised using NormalizeData (scale factor = 10,000 by default) and were used to run FindMarkers for identifying additional cluster marker genes for each annotated cluster.

#### Pseudobulked Differential Gene Expression Analysis and Pathway Analysis

The normalised count matrix and the metadata were extracted from the Seurat object. Counts were aggregated per sample (i.e. donor and day) for each cluster. Since we conducted a paired-wise comparison (e.g. day 0 vs day 16) only clusters present in both samples were included in the analysis. Significant genes were determined using the quasi-likelihood (QL) F-test with an FDR< 0.05 using edgeR ^63^. Aggregated counts were used for visualising changes in specific DEGs using ComplexHeatmap.

#### TCRA/B clonal analysis

Annotated TCR sequences were integrated with the single-cell transcriptomics data using the combineTCR and combineExpression functions from scRepertoire (version 2.0.0). The CDR3 amino acid sequences were used to call and quantify clones. Only cells with single and complete paired TCRA and TCRB chain were included. The R package circlize (v0.4.16) and clonalOverlap function of scRepertoire were used to generate the circos and heatmap plots of clonal overlap, respectively. For Simpson’s diversity index calculations, data were down-sampled to 1,000 cells to adjust for cell count variations between samples.

### Statistical analysis

Non-parametric testing was performed for all analysis. Continuous data for all cellular responses were compared between groups using Mann-Whitney U test, or in paired samples with Wilxcox paired sign-rank test, or correlated with antibody score as a continuous variable with Spearman’s correlations. Statistical comparisons were adjusted for multiple comparisons where indicated. All analyses were performed in R (version 4.4.2). Graphical outputs were made in ggplot2 (version 3.5.1) and ggpubr (version 0.6.0). No sample size calculation was performed, instead all available participant data was included. Due to sample size restrictions, no subgroup analysis was performed. Data generation was performed with blinding to participant demographic data.

### Data and code availability

All data generated or analysed during this study are included in this article and supplementary information files or deposited online as follows. The raw sequencing data and processed Cell Ranger outputs used in this study have been deposited in the NCBI data base under accession code GSE272939 https://www.ncbi.nlm.nih.gov/geo/query/acc.cgi?acc=GSE272939 and GSE253661 https://www.ncbi.nlm.nih.gov/geo/query/acc.cgi?acc=GSE253661.

The processed single cell RNA sequencing data are available in the Zenodo, https://doi.org/10.5281/zenodo.14847353.

Code used to analyse scRNAseq data is available at https://github.com/MichelleBoyle/CCR7-expression-defines-distinct-pTfh1-subsets-involved-in-malaria-immunity.git

Flow cytometry files will be made freely available on publication in an online repository.

## Author contributions

MSFS, ND, RM, DA, JRL and JAC generated data, supervised by JGB, CE, AH and MJB MSFS, DAO, ZP analysed data

JM provided clinical samples used in this study AH and MJB conceptualised study

MSFS and MJB led manuscript writing.

All authors contributed to and approved manuscript

## Acknowledgements

Burnet Institute, University of Melbourne and QIMR-Berghofer acknowledge the traditional custodians of the lands where they are located, the Boonwurrung and Wurundjeri Woi-wurrong people of the Kulin Nation, and the Turrbal and Jagera people.

RBC and human serum were provided by the Australian Red Cross Blood Bank (Melbourne and Brisbane). We thank the participants involved in the CHMI studies and all study clinicians and support staff at QPharm.

This work was supported by the National Health and Medical Research Council of Australia (program grant 1132975 to J.S.M and C.R.E.); Senior Research Fellowships C.R.E. (1154265), Career Development Award 1141278, Project Grant 1125656, and Ideas Grant 1181932 to MJB); the CSL

Centenary Fellowship and the Snow Medical Foundation Fellowship 2022/SF167 to M.J.B; the Jim and Margaret Beever Fellowship to J.A.C. The Burnet Institute is supported by the NHMRC for Independent Research Institutes Infrastructure Support Scheme and the Victorian State Government Operational Infrastructure Support. Medicines for Malaria Venture funded the parent CHMI studies.

