## Supplementary Tables and Figures for "Functionally Distinct pTfh1 Subsets with Roles in Malaria-Specific Immunity"

**Supplementary Figures and Tables**

**Supplementary Tables.**

***Supplementary Table S1:*** Cluster markers for pTfh clusters in healthy donors *attached as excel*

***Supplementary Table S2:*** DEGs between Tfh1_cyto and Tfh1_CCR7 clusters *attached as excel*

***Supplementary Table S3: Phenotyping panel***

| Target | Fluorochrome | Clone | Manufacturer | Dilution |
|---|---|---|---|---|
| LAG-3 | BUV661 | 3DS223H | Invitrogen | 1 in 25 |
| CD49b | FITC | eBioY418 | Invitrogen | 1 in 50 |
| CD25 | BUV395 | 2A3 | BD Biosciences | 1 in 100 |
| CXCR5 | BUV615 | MU5UBEE | ThermoFisher | 1 in 50 |
| CXCR3 | BV421 | 1C6/CXCR3 | BD Biosciences | 1 in 25 |
| CCR6 | BV605 | 11A9 | BD Biosciences | 1 in 50 |
| OX40 | BV650 | ACT35 | BD Biosciences | 1 in 50 |
| CCR7 | BV711 | 150503 | BD Biosciences | 1 in 50 |
| CD3 | BV786 | UCHT1 | BD Biosciences | 1 in 500 |
| CD4 | Spark blue 574 | SK3 | Biolegend | 1 in 100 |
| CD69 | PerCP-eFluor 710 | FN50 | ThermoFisher | 1 in 100 |
| TIM-3 | PE-CF594 | 7D3 | BD Biosciences | 1 in 50 |
| PD-1 | PE-Fire 640 | EH12.2H7 | Biolegend | 1 in 50 |
| TIGIT | PE-Fire 810 | A151513G | Biolegend | 1 in 100 |
| CD45RA | redFluor 710 | HI100 | Cytek Biosciences | 1 in 400 |
| ICOS | APC-H7 | DX29 | BD Biosciences | 1 in 100 |
| CD38 | APC-Fire 810 | HIT2 | Biolegend | 1 in 200 |
| **Intracellular:** | | | | |
| CTLA-4 | BUV737 | 14D3 | ThermoFisher | 1 in 50 |
| Ki-67 | BUV805 | B56 | BD Biosciences | 1 in 50 |
| Granzyme B | BV510 | GB11 | BD Biosciences | 1 in 800 |
| Granzyme K | RB780 | G3H69 | BD Biosciences | 1 in 50 |
| NKG7 | PE | E6S2A | Cell Signalling Technology | 1 in 2000 |
| c-Maf | PECy7 | sym0F1 | ThermoFisher | 1 in 400 |
| FOXP3 | RB613 | 259D/C7 | BD Biosciences | 1 in 100 |
| SAP | eFluor660 | XLP-1D12 | ThermoFisher | 1 in 100 |

***Supplementary Table S4: PMA/Io panel***

| Target | Fluorochrome | Clone | Manufacturer | Dilution |
|---|---|---|---|---|
| CD8 | BUV496 | RPA-T8 | BD Biosciences | 1 in 100 |
| CD45RA | BUV563 | HI100 | BD Biosciences | 1 in 400 |
| CD3 | BUV805 | SK7 | BD Biosciences | 1 in 50 |
| CD14/CD19 | BV510 | M5E2 / SJ25C1 | Biolegend | 1 in 100 |
| CCR4 | BV605 | L291H4 | Biolegend | 1 in 100 |
| CCR6 | BV650 | 11A9 | BD Biosciences | 1 in 50 |
| CXCR5 | BV711 | J252D4 | Biolegend | 1 in 50 |
| CD4 | BV785 | OKT4 | Biolegend | 1 in 50 |
| CXCR3 | PE-CF594 | 1C6/CXCR3 | BD Biosciences | 1 in 50 |
| PD-1 | PECy7 | EH12.1 | BD Biosciences | 1 in 50 |
| Vd2 | APC-Fire | B6 | Biolegend | 1 in 50 |
| **Intracellular:** | | | | |
| IFNy | BUV395 | B27 | BD Biosciences | 1 in 50 |
| IL-10 | BV421 | JES3-9D7 | Biolegend | 1 in 10 |
| TNFa | BV750 | MAb11 | BD Biosciences | 1 in 100 |
| IL-17a | FITC | BL168 | Biolegend | 1 in 25 |
| IL-21 | PE | 3A3-N2.1 | BD Biosciences | 1 in 10 |
| IL-4 | APC | MP4-25D2 | Biolegend | 1 in 10 |

***Supplementary Table S5***: Genes used in input for scType prediction *attached as excel*

***Supplementary Table S6:*** Cluster markers for predicted annotated cell clusters in CHMI data set *attached as excel*

***Supplementary Table S7:*** DEGs in each Tfh subset during malaria infection identified by edgeR pseudobulk *attached as excel*

***Supplementary Table S8:*** Antibody panel for sort purification of Tfh cells for scRNAseq of healthy samples

| Target | Fluorochrome | Clone | Manufacturer | Dilution |
|---|---|---|---|---|
| CD45RA | BB515 | HI100 | BD Biosciences | 1 in 400 |
| CD4 | PerCpCy5.5 | OKT4 | Biolegend | 1 in 50 |
| CXCR3 | PE-CF594 | 1C6/CXCR3 | BD Biosciences | 1 in 50 |
| PD-1 | PECy7 | EH12.1 | BD Biosciences | 1 in 50 |
| CD3 | AF700 | OKT3 | Biolegend | 1 in 50 |
| | Sytox Blue | | | |
| CCR6 | BV650 | 11A9 | BD Biosciences | 1 in 50 |
| CXCR5 | BV711 | J252D4 | Biolegend | 1 in 50 |

***Supplementary Table S9:*** Antibody panel for sort purification of Tfh cells for scRNAseq of CHMI samples

| Target | Fluorochrome | Clone | Manufacturer | Dilution |
|---|---|---|---|---|
| **Stain 1:** | | | | |
| TCR gD | FITC | B1 | Biolegend | 3 in 25 |
| CXCR5 | BV711 | J252D4 | Biolegend | 2 in 25 |
| **Stain 2:** | | | | |
| CD4 | PerCpCy5.5 | OKT4 | Biolegend | 1 in 50 |
| CD19 | PE | HIB19 | BD Biosciences | 1 in 100 |
| PD-1 | PECy7 | EH12.1 | BD Biosciences | 1 in 50 |
| Vd2 | APC | B6 | Biolegend | 1 in 50 |
| CD3 | AF700 | OKT3 | Biolegend | 1 in 50 |
| | Sytox Blue | | | |
| CD56 | BV510 | HCD56 | Biolegend | 1 in 100 |
| CD45RA | BV570 | HI100 | Biolegend | 1 in 400 |
| HLA-DR | BV785 | L243 | Biolegend | 1 in 200 |

### Supplementary Figures

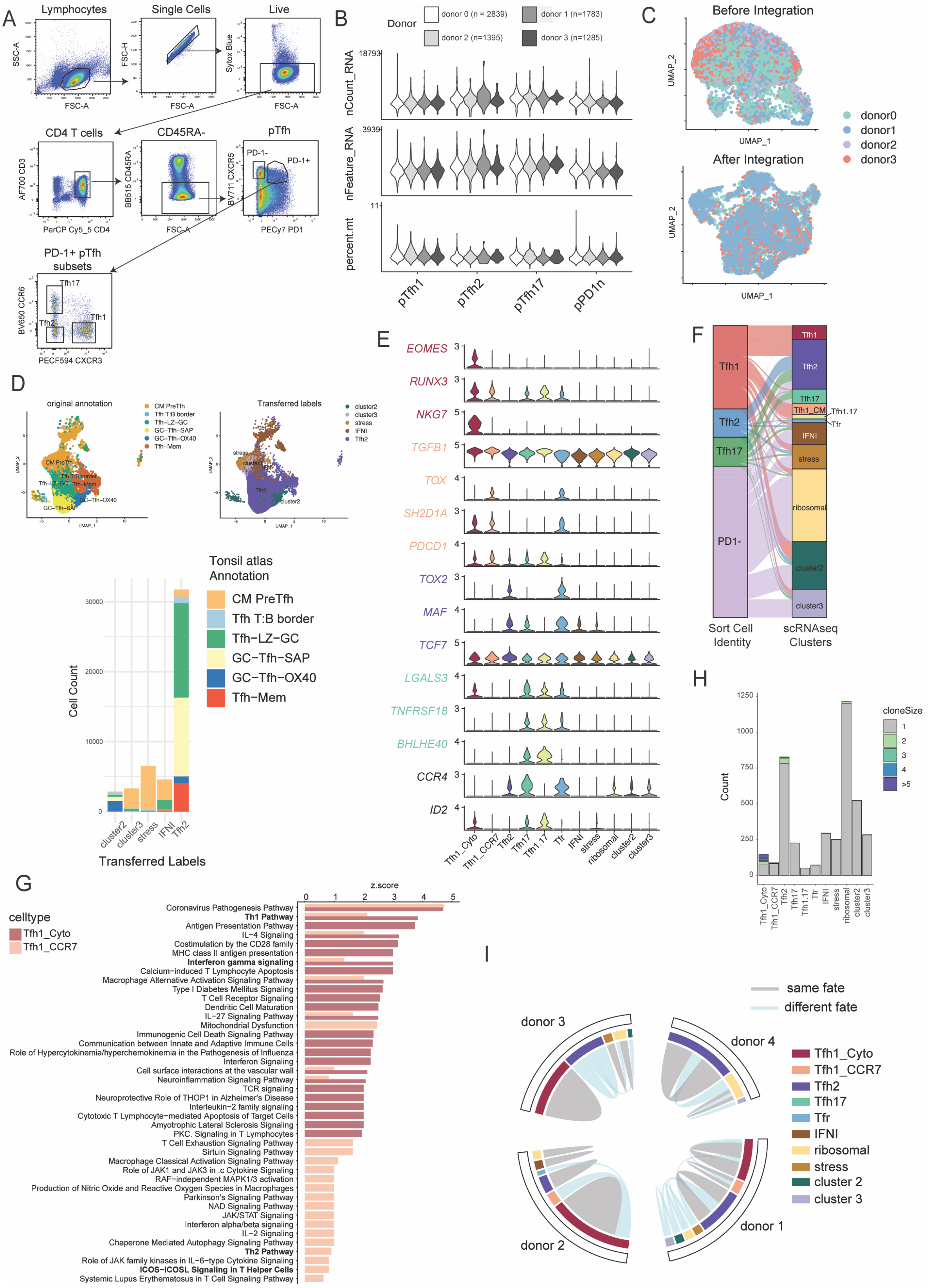

***Supplementary Figure 1. Single cell RNA sequencing of pTfh cells from healthy donors.***

***A)** Gating strategy for sorting pTfh cell populations scRNAseq. Non-naïve CD4 T cells were identified as CD3+/CD4+, CD45RA- cells. Tfh1 (CXCR3+CCR6-), Tfh2 (CXCR3-CCR6-), Tfh17(CXCR3-*

*CCR6+) were sorted from CXCR5+PD1+ cells, and a resting Tfh cell population from CXCR5+PD1- cells.* ***B)*** *QC of cells from each donor and subset, nCount_RNA, nFeature_RNA and percent mitochondrial content is shown.* ***C)*** *UMAP of data before and after integration for donor.* ***D)*** *Barchart showing the number of cells in each predicted cell cluster grouped by their original cell annotation from the human tonsil atlas.* ***E)*** *Expression of genes with various roles in Tfh and CD4 T cell development across different Tfh cell clusters.* ***F)*** *Relationship between cell sorted identity and transitional cell cluster identify.* ***G)*** *Pathway analysis of top 25 pathways curated from cluster marker genes of Tfh1_CCR7 and Tfh1_cytotoxic clusters.* ***H.*** *Frequency of individual clones that occurred with clonal size of 1, 2, 3, 4, >5 in each subset.* ***I.*** *Circos plot of clonal overlap between subsets. No clones were shared across donors. Expanded clonotypes (n$^{3}$2) with only same fates (exist within the same subset), or different fates are shown.*

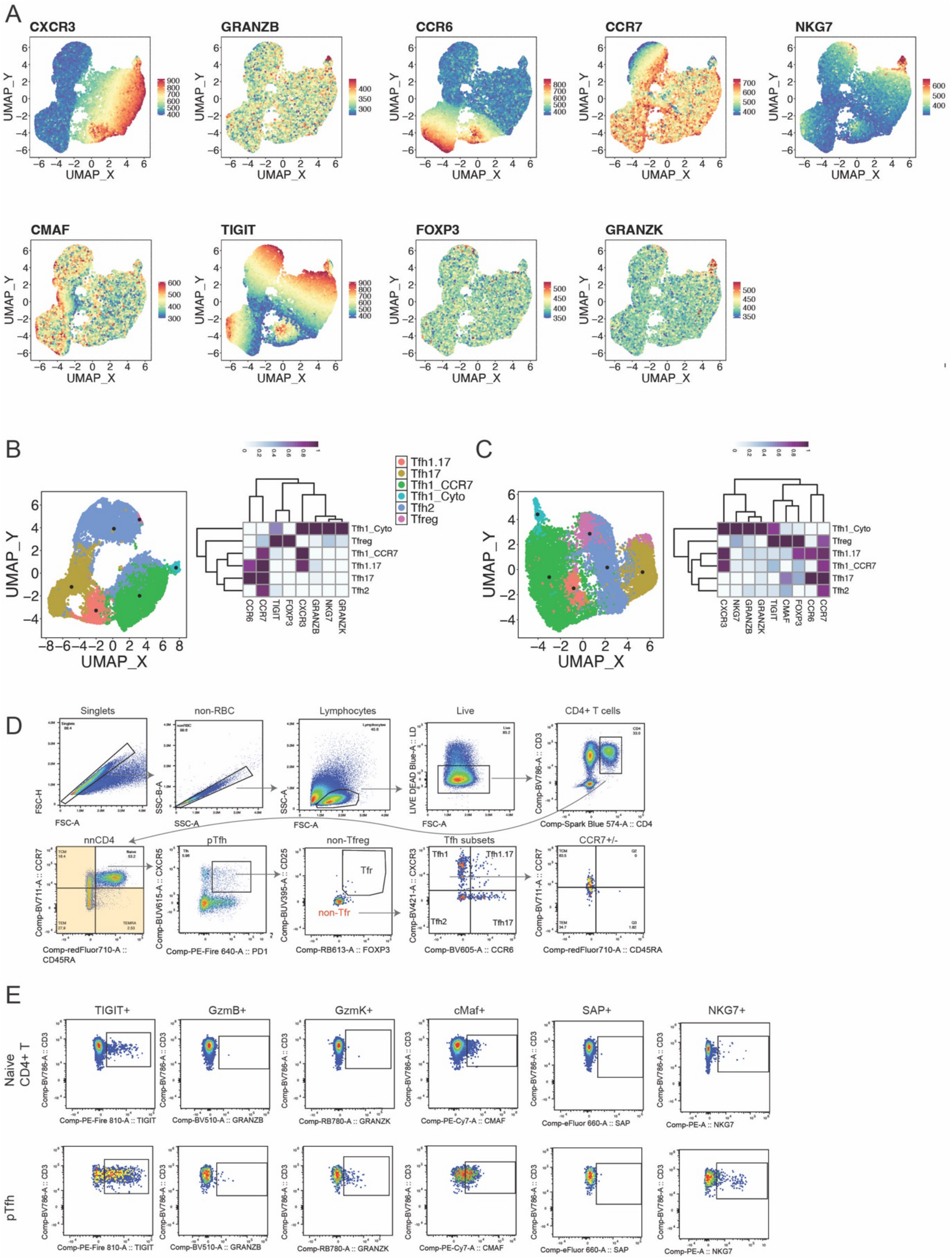

**_Supplementary Figure 2: Phenotypic diversity within pTfh subsets based on CCR7 expression._**

***A.*** *Expression of each marker used in unbiased analysis (see Figure 2).* ***B/C****. UMAP of unbias analysis of a single experiment (n=6 for E and n=Y for 6).* ***D.*** *Gating strategy for identify pTfh cells. pTfh cells were identified as CXCR5+PD1+ non-naïve CD4 T cells. Within pTfh cells, Tfr cells were identified as FoxP3+/$CD127^{lo}$ cells, pTfh subsets identified based on CXCR3 and CCR6, and CCR7 expression analysed.* ***E.*** *Marker expression in naïve CD4 T cells and pTfh cells.*

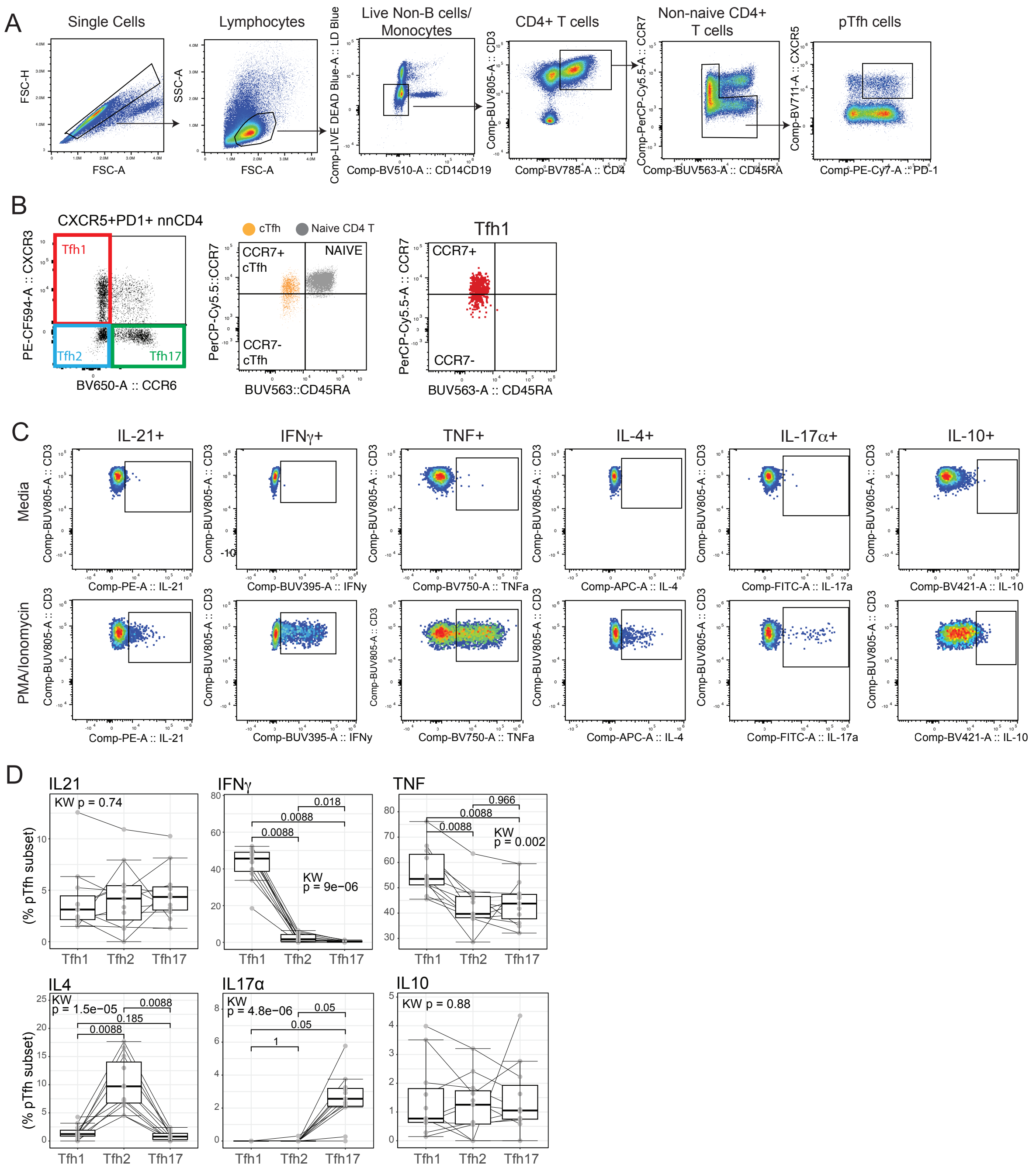

***Supplementary Figure 3: Gating strategy for pTfh cells and cytokine expression following PMA/Io stimulation.***

*PBMCs were stimulated with PMA/Ionomycin to assess cytokine production.* ***A)*** *pTfh cells were identified as CXCR5+PD1+ CD4 T cells.* ***B)*** *A gradient of CCR7 could be detected within pTfh1 cell susbsets, and pTfh1 clusters can be divided into $CCR7^{pos}$ and $CCR7^{neg}$ cells.* ***C)*** *Representative data of cytokine expression in pTfh cells in unstim (media) or PMA/Ionomycin stimulated cells.* ***D)*** *pTfh cells subsets were identified by CXCR3 and CCR6 expression as pTfh1 (CXCR3+CCR6+), pTfh2 (CXCR3-CCR6-) and pTfh17 (CXCR3-CCR6+) cells (grouping $CCR7^{pos}$ and $CCR7^{neg}$ Tfh1 cells). P is the paired Wilcoxon signed-rank test between groups, adjusted for multiple comparisons with Holm's FDR, and Kruskal Wallis group comparison test.*

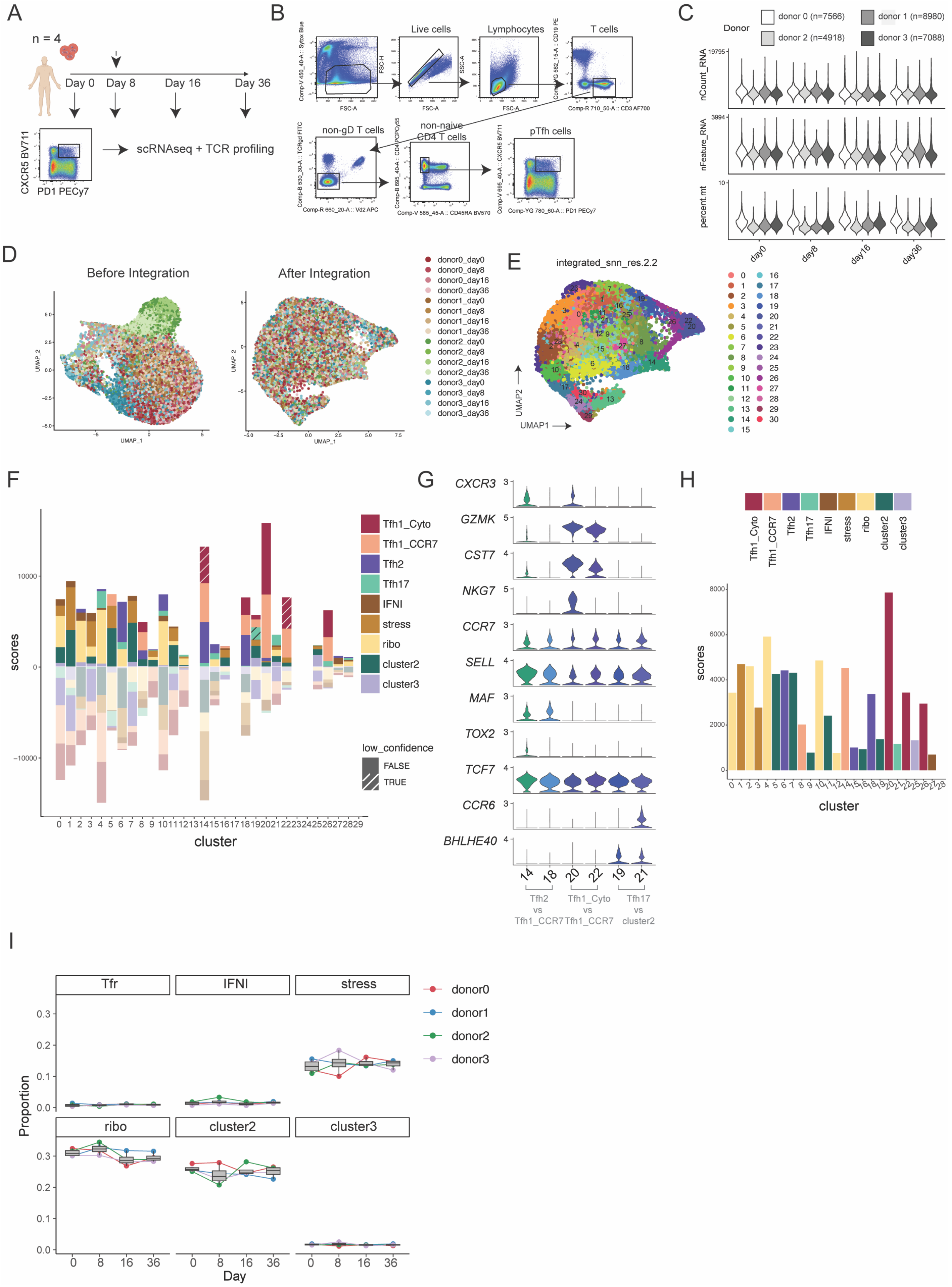

**Supplementary Fig 4 scRNAseq analysis of Tfh cells in CHMI**

***A)*** *scRNAseq experimental set up for Tfh cells during controlled human malaria infection (CHMI). Tfh cells (CXCR5+PD1+) identified within non-naïve CD4 T cells were sorted from four individuals at day 0, 8, 16 and end-of-study during CHMI. Individuals were treated at day 8 (indicated by dotted* *arrow).* ***B)*** *Gating strategy for Tfh cell sorting for scRNAseq.* ***C)*** *QC of data from each donor across time points nCount_RNA, nFeature_RNA and percent mitochondrial content is shown.* ***D)*** *Data before and after integration based on donor and CHMI day.* ***E)*** *Over clustering of data prior to scType label transferer.* ***F)*** *scType was used to label transfer cell signatures (top 20 up and down regulated genes) for each cell subset identified in healthy 'map' data onto over-clustered CHMI data. Each cell cluster was scored for each reference signature (left), and the majority score (right) used to collapse clusters into cell subsets.* ***G) Average expression of*** *Tfh cell cluster markers in each subset for manual adjustment of "low-confidence" predicted cluster identities.* ***H).*** *Final predicted label for each cell cluster.* ***I****) Proportion of other Tfh cell subsets identified over time.*

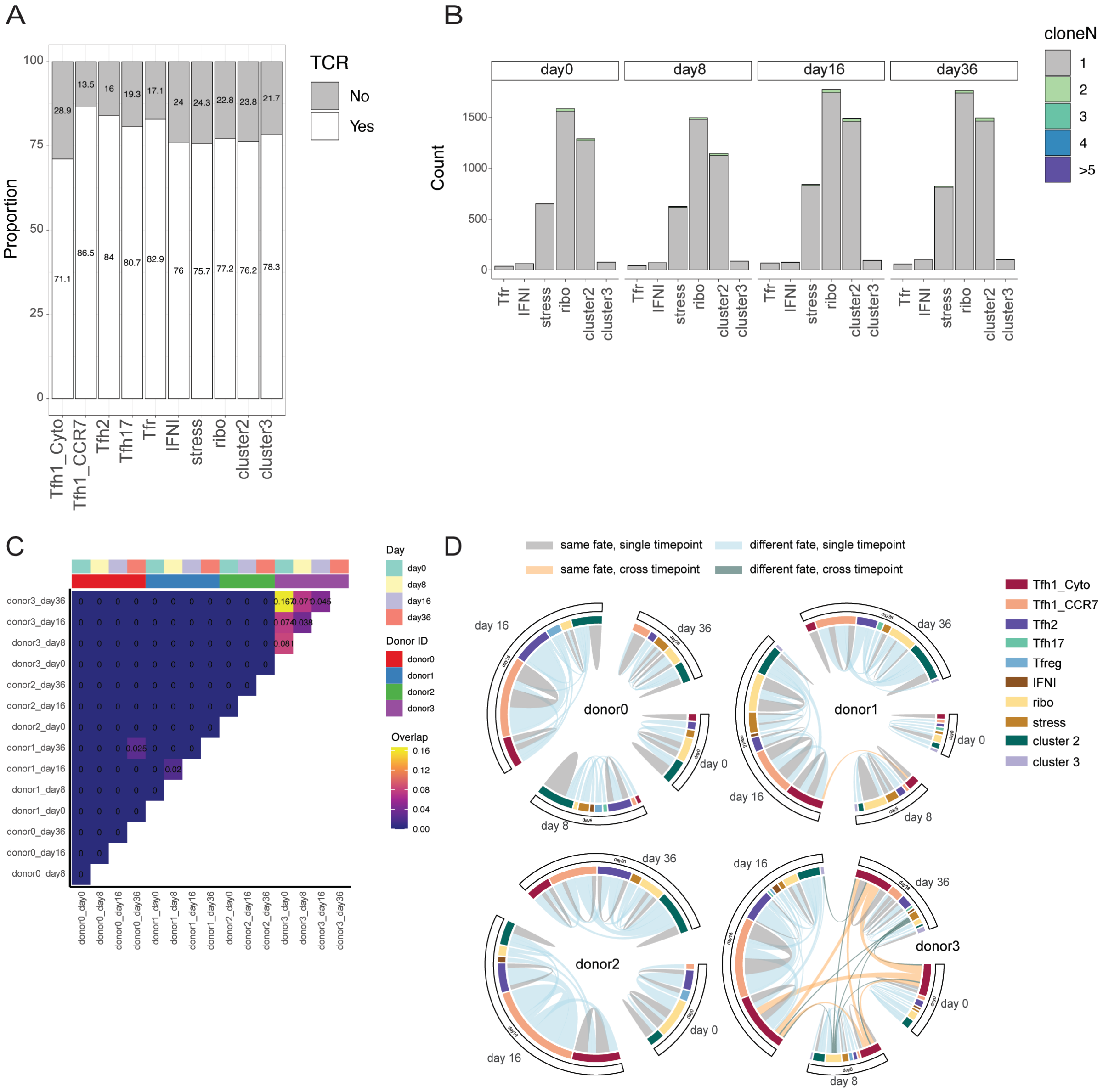

***Supplementary Figure 5: TCR analysis of Tfh cells in CHMI***

***A)*** *Proportion of cells with TCR captured across each subset.* ***B)*** *Clonal size of other Tfh cell subsets identified over time.* ***C)*** *Clonal overlap between donor and day.* ***D)*** *Circos plot indicating clonal overlap between cell clusters and day for each donor.*

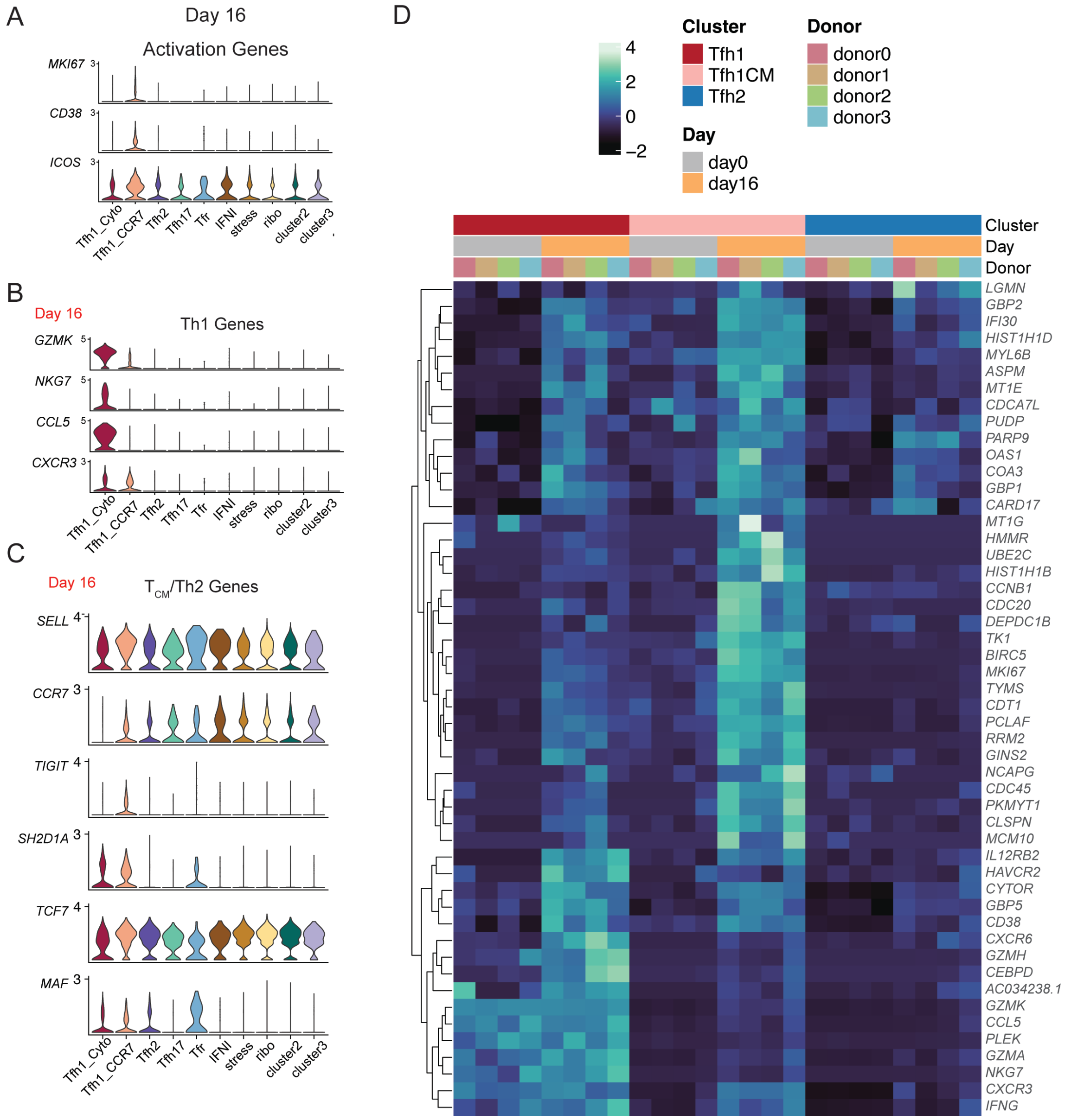

***Supplementary Figure 6: Tfh cell profiles at day 16 following CHMI***

*DEGs were identified in each subsets comparing day 0 to subsequent time points during infection.* ***A)*** *Expression of activation genes for all clusters at Day 16.* ***B-C)*** *Despite upregulation of genes associated with inflammation, expression profiles were still different between Tfh1_cyto and Tfh1_CCR7 subsets. Tfh cell subsets maintained expression profiles of identifying cluster markers.* ***D)*** *Heatmap showing the top 50 unique genes (curated from top 20 upregulated genes of Tfh1_Cyto, Tfh1_CCR7 and Tfh2 clusters) upregulated at Day 16 for Tfh1_Cyto, Tfh1_CCR7 and Tfh2 clusters relative to Day 0.*

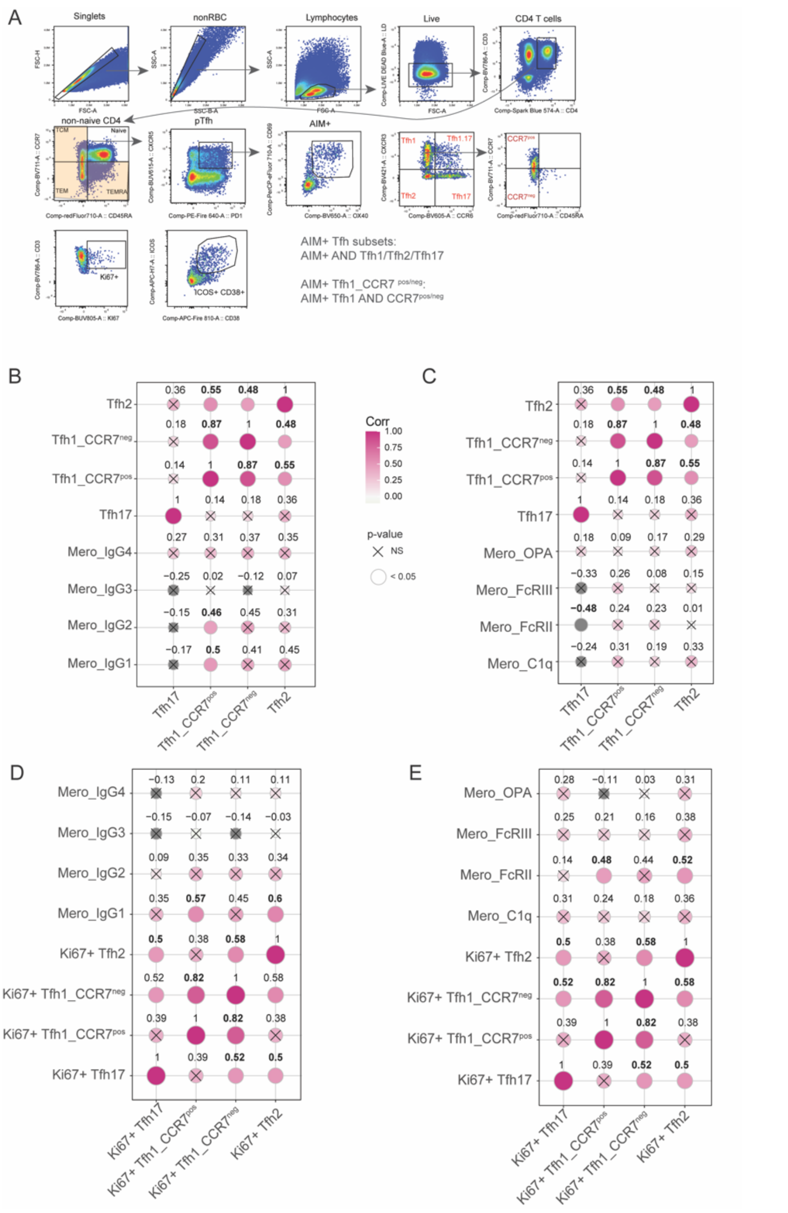

***Supplementary Figure 7: Validating Tfh cell profiles in CHMI***

***A)** Gating strategy for identification of malaria specific pTfh cells in individuals undergoing CHMI using Activation Induced Marker assay. **B-E)** Correlation matrix between malaria-specific Tfh subsets **(B-C)** or Ki67 expression of Tfh subsets **(D-E)** at day 36 with either IgG subclasses to merozoite **(B/D)** or functional capacity to fix complement (C1q), bind dimeric FcγRIIa or FcγRIIIa (surrogates for IgG antibody capacity to crosslink cellular receptors), and promote opsonic phagocytosis (OPA) **(C/E)**. **B-E.** Spearman's rho and p-value are indicated.*
